# JMnorm: a novel Joint Multi-feature normalization method for integrative and comparative epigenomics

**DOI:** 10.1101/2023.06.14.545004

**Authors:** Guanjue Xiang, Yuchun Guo, David Bumcrot, Alla Sigova

## Abstract

Combinatorial patterns of epigenetic features reflect transcriptional states and functions of genomic regions. While many epigenetic features have correlated relationships, most existing data normalization approaches analyze each feature independently. Such strategies may distort relationships between functionally correlated epigenetic features and hinder biological interpretation. We present a novel approach named JMnorm that simultaneously normalizes multiple epigenetic features across cell types, species, and experimental conditions by leveraging information from partially correlated epigenetic features. We demonstrate that JMnorm-normalized data can better preserve cross-epigenetic-feature correlations across different cell types and enhance consistency between biological replicates than data normalized by other methods. Additionally, we show that JMnorm-normalized data can consistently improve the performance of various downstream analyses, which include candidate cis-regulatory element clustering, cross-cell-type gene expression prediction, detection of transcription factor binding and changes upon perturbations. These findings suggest that JMnorm effectively minimizes technical noise while preserving true biologically significant relationships between epigenetic datasets. We anticipate that JMnorm will enhance integrative and comparative epigenomics.

**GRAPHICAL ABSTRACT:** JMnorm can jointly normalize multiple epigenetic features between the target sample and the reference.

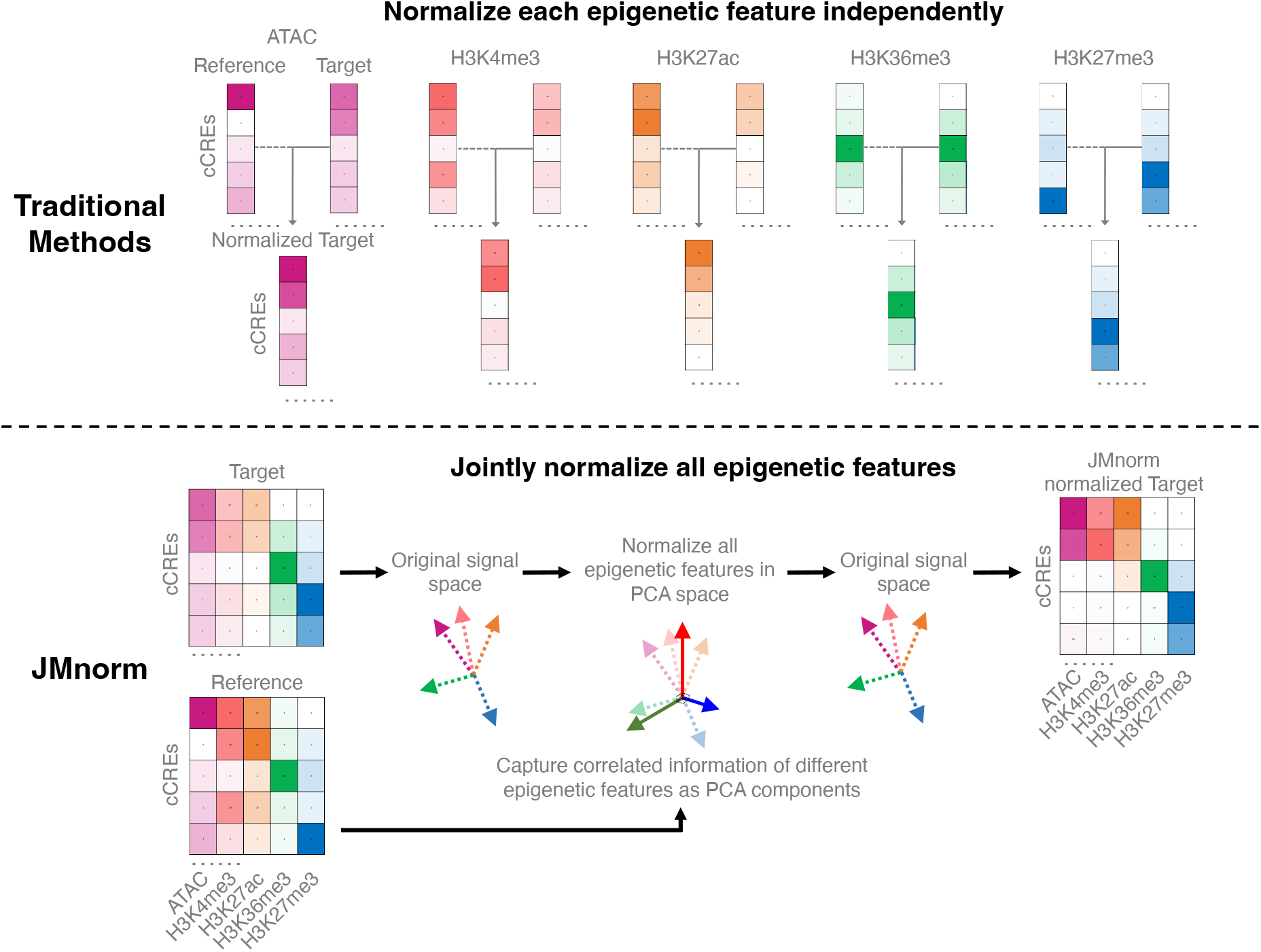

## INTRODUCTION

Epigenetic features, including DNA accessibility and posttranslational histone modifications, are thought to accurately reflect transcriptional states and help infer mechanistic insights about the regulation of gene expression in cell-type-specific contexts. Development of high-throughput sequencing techniques for genome interrogation led to the generation of hundreds of epigenetic datasets in different cell types and under various physiological conditions. Many large-scale data consortiums, such as the Encyclopedia of DNA Elements (ENCODE) and the ValIdated Systematic IntegratiON of hematopoietic epigenomes (VISION) projects, have utilized epigenetic features to identify candidate cis-regulatory elements (cCREs) (1–4). Follow-up studies characterized functional dynamics of epigenetic features across different cell types and conditions to elucidate their effects on transcriptional regulation (5–10). Increased sensitivity of the high-throughput sequencing methods results in amplified technical noise that can hinder the ability to extract biologically meaningful information. Therefore, to precisely quantify and compare epigenetic features across cell types, species, and experimental conditions, it is essential to develop robust epigenomic data normalization techniques to mitigate technical biases (11).

Many normalization methods have been developed for comparative analyses of high-throughput sequencing data (Supplementary Table 1). The two most widely used and easily implementable normalization methods are total library size normalization (TSnorm) and quantile normalization (QTnorm). TSnorm scales the signal of datasets to be compared based on the ratio of their total library sizes (12, 13), whereas QTnorm transforms and equalizes signal distribution of each dataset relative to a reference distribution (14). The advantage of TSnorm is in its simplicity as it assumes that the only source of technical variation between datasets lies in differences in their sequencing read depths. QTnorm, on the other hand, can effectively remove complex technical biases by assuming different datasets share not only the same global mean but also the same global distribution. These assumptions hold true for certain types of high-throughput sequencing data such as bulk cell RNA-seq, where the majority of true biological signals are similar in different data sets. However, they are likely incorrect for epigenetic datasets with substantial variability in the number of peaks and signal intensities across different cell types (1, 2) or experimental conditions (15), especially for epigenetic features with prominent signals at the cell-type-specific enhancers (1, 2). Consequently, in comparative analyses of data with unequal number of biological peaks between cell types or conditions, both normalization methods tend to generate false positive or negative peak signals. Moreover, when comparing datasets produced under different treatment conditions inducing global epigenomic effects, QTnorm often distorts normalized signals relative to the true signals.

Other more specialized normalization methods, for example MAnorm and S3norm and their latest versions (16–19), are more adept for analysis of epigenetic datasets. These methods aim to eliminate biases due to differences in both sequencing depths and signal-to-noise ratios, which often arise from factors such as variations in antibody efficiency during ChIP-seq experiments (20). They utilize information from shared peak regions without/with common background regions and assume that the true signals in these regions remain consistent across different datasets and thereby, can serve as reliable anchors for data normalization. However, the simple scaling factors or transformation models employed by these methods might not be adequate for addressing all technical biases, especially when they exhibit complex patterns, which are often observed in studies integrating datasets from multiple sources (3, 21). Lastly, some methods can effectively model diverse signal distributions of epigenetic data by converting signals into ranks. However, these methods result in a loss of quantitative information (22).

Another significant drawback of existing approaches is that they analyze each epigenetic feature independently. These approaches may distort relationships between functionally correlated epigenetic features and hinder biological interpretation. Combinatorial patterns of multiple epigenetic features, known as epigenetic states, have been widely used for functional annotation of cCREs in different cell types, species, and experimental conditions (3, 23–28). Recent studies have shown that, while regulatory regions with specific epigenetic states may vary based on cellular and experimental contexts, cross-feature combinatorial patterns remain relatively conserved (4). Therefore, a normalization method that utilizes the information from functionally correlated epigenetic features could yield more accurate and biologically relevant post-normalization signals, enabling more meaningful comparison and integration of epigenetic data across conditions.

Here, we present a novel approach called <u>J</u>oint <u>M</u>ulti-feature <u>norm</u>alization (JMnorm) for simultaneous normalization of multiple epigenetic features across cell types, species, and experimental conditions by leveraging information from functionally correlated features. We demonstrate JMnorm’s superior performance in preserving cross-feature correlations and improving consistency between biological replicates, as well as its better versatility and utility for various genomic applications relative to other methods. Additionally, JMnorm can increase consistency between normalized epigenetic features and orthogonal datasets, which we robustly validated across diverse types of epigenetic features, including transcription factor (TF) binding ChIP-seq and DNase-seq data.

## MATERIALS AND METHODS

### Data collection and preprocessing

We obtained epigenetic datasets primarily from two databases: the VISION (3, 4, 21, 29–32) and the EpiMAP repository (33, 34). For VISION datasets, we downloaded bigWig files containing average read counts for seven chromatin features (H3K4me3, H3K4me1, H3K27ac, H3K36me3, H3K27me3, H3K9me3, ATAC-seq) from 9 human and 16 mouse hematopoietic cell types. For EpiMAP datasets, we downloaded bigWig files containing average −log10(p-value) signal tracks for seven chromatin features (H3K4me3, H3K4me1, H3K27ac, H3K36me3, H3K27me3, CTCF ChIP-seq, and DNase-seq) from 24 human cell type groups. Since EpiMAP signal tracks were originally mapped to the hg19 reference genome, we used the CrossMap package (35) with default settings to lift over these files to the hg38 reference genome. All other datasets used in this study were mapped to hg38. RNA-seq data for different cell types were also downloaded from the VISION project (log2TPM, quantile normalized) and the EpiMAP repository (log2FPKM, quantile normalized). The topologically associating domains (TAD) boundaries were downloaded from the VISION project website under the “Hi-C” tab (36), and YY1 peaks (bed format) were downloaded from the Cistrome DB (37–39). The links to the downloaded files and the Cistrome DB sample ID list of the downloaded YY1 peak files are provided in Supplementary Table 1. The DNase-seq data to obtain the number of DNase I Hypersensitive Sites (DHSs) in different cell types were downloaded from the Meuleman 2020 study (40, 41). To prepare data for normalization and downstream analyses, we first acquired the epigenomic signal matrices. This was accomplished by calculating the average signals for each chromatin feature at each ENCODE cCRE region (2) in every biological replicate of each cell type using the bigWigAverageOverBed script from the UCSC Genome Browser utilities (42). Subsequently, we generated a reference signal matrix by combining the average signal for each chromatin feature and used the matrix for all datasets to normalize against. It is important to note that not all cell types had datasets for all epigenetic features. The details and links to all available datasets in both species are provided in Supplementary Table 1.

### Generation of reference signal matrix

To normalize data across multiple cell types, we used the average signal as reference for all cell types to normalize against in our study. For each epigenetic feature, we first collected all datasets for that feature in all cell types, and then computed the average signals at each cCRE region across these cell types. The average signal vector for all cCREs was used as a reference signal vector for the specific feature. This process was performed for all features, and the resulting outputs were combined to generate the reference signal matrix.

To improve the cross-feature comparability and interpretability, we further equalized sequencing depths and signal-to-noise ratios of different features in the reference matrix using S3norm (17). Instead of using S3norm’s default mode, which learns an exponential transformation model from common peak regions and common background regions of two signal vectors, we employed its cross-feature normalization mode. This mode leverages peak regions and background regions of each signal vector to learn a transformation model. The reason for this choice is that when normalizing two signal vectors of different epigenetic features, the assumption that common peak regions represent housekeeping epigenetic events with similar signal levels is not valid, particularly for features with opposing functions such as H3K27ac and H3K9me3. Specifically, we used the ratio between the top mean signal of 99% quantile of cCREs and the overall mean of all cCREs for each feature to learn the S3norm transformation model. After the normalization, the sequencing depth and the signal-to-noise ratios are equalized across all features in the reference matrix. This reference signal matrix is then used as the reference signal matrix for all cell types to normalize against in downstream analyses.

### The details about JMnorm

JMnorm consists of four key steps (Figure 1). In the first step, it transforms the signal matrix into principal component analysis (PCA) space to consolidate correlated components across the signal vectors of different epigenetic features into the same PCA dimension. To achieve this, JMnorm learns a PCA transformation model from the reference signal matrix and then applies it to transform the target signal matrices. This ensures the signal matrices of all target cell types can be transferred to the same reference PCA spaces. Moreover, the PCA model captures the cross-feature correlation information from the reference signal matrix, which is used and preserved throughout the subsequent steps.

**Figure 1.**
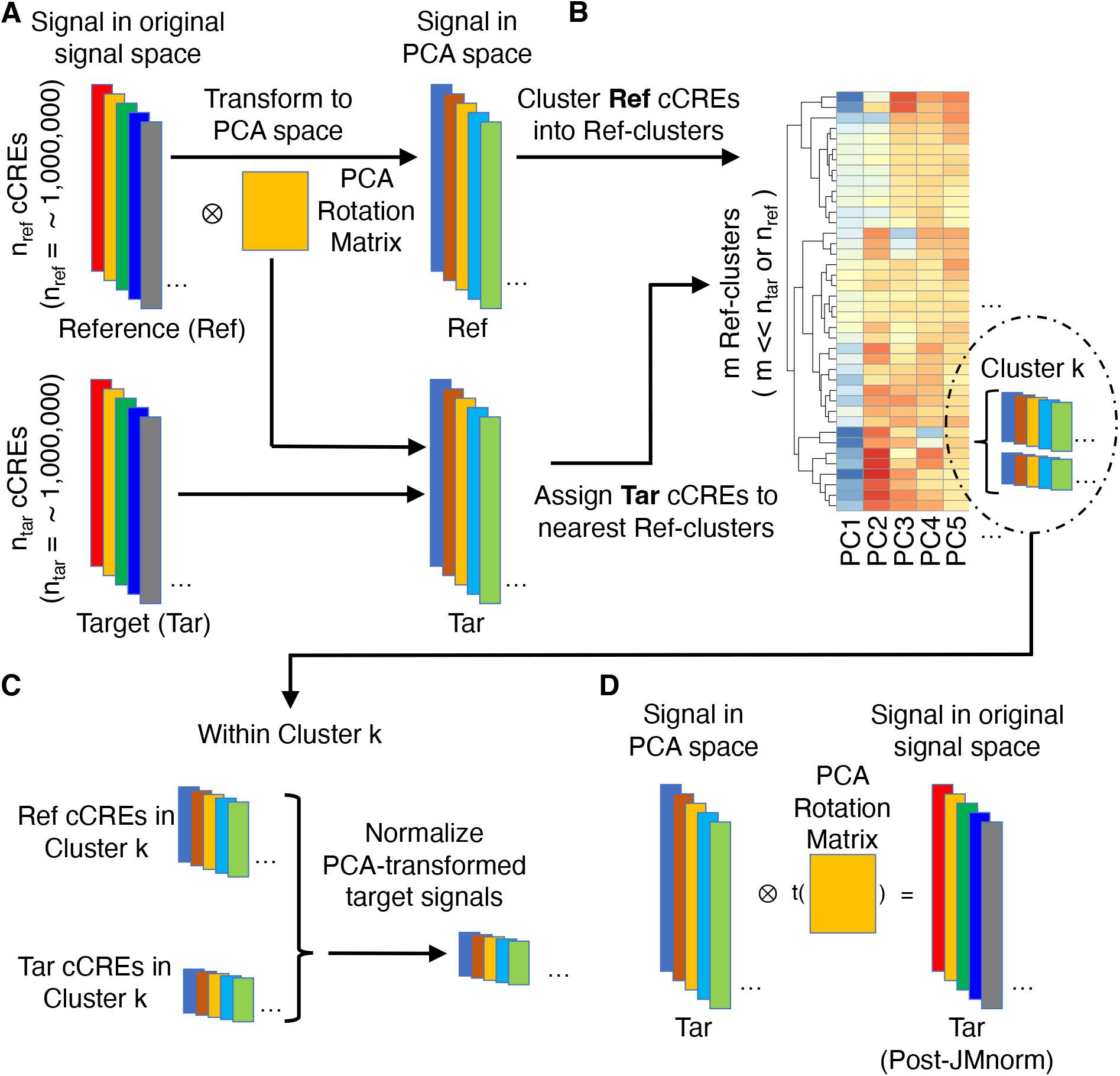
An overview of the four key steps in the JMnorm normalization procedure. (A) Step 1: orthogonal transformation. The correlated components of various epigenetic signals are transformed into mutually independent high-dimensional PCA dimensions. Each colored block on the left represents the signal vector of all epigenetic features at the n_ref_ or n_tar_ cCRE regions in reference or target samples, respectively. Colored blocks on the right denote corresponding transformed PCA epigenetic signal matrices for reference and target samples. The yellow box in the middle represents the PCA rotation matrix learned from the reference signal matrix. (B) Step 2: cCRE clustering. Reference cCRE clusters are generated based on the reference data in the PCA space with the average signal reference matrix shown as a heatmap. Target cCREs are assigned to reference clusters according to the Euclidean distances between the signal vector of the target cCRE and the average signal vectors of reference clusters in the PCA space. Within each cluster, the number of cCREs, shown as colored blocks within the insert, may vary between the reference and target samples. (C) Step 3: within-cluster normalization. Target signal matrix is normalized against the reference matrix using within-cluster quantile normalization as shown for Cluster k. (D) Step4: reconstruction of the JMnorm-normalized target signal matrix in the original signal space. The yellow box in the middle indicates the transposed PCA rotation matrix learned in the first step (panel A).

Previous studies have demonstrated that there are reproducible combinatorial patterns across various chromatin features in different cell types or species, commonly referred to as epigenetic states, which play a role in transcriptional regulation. In the second step, we leveraged this prior knowledge by first clustering the cCREs into distinct groups based on the reference signal matrix (reference clusters). The reference clusters were generated using K-means clustering based on the reference signal matrix in PCA space. The number of reference clusters (K) was automatically determined by performing hierarchical clustering on a subset of data (20,000 cCREs by default), followed by the DynamicTreeCut method (cutreeDynamic function with the following parameter settings: method=‘hybrid’ and deepSplit=2) (43). The resulting number of DynamicTreeCut clusters was then utilized as the K for K-means clustering on the entire reference signal matrix. Under the assumption that these reference clusters capture the conserved cross-feature patterns of distinct cCRE groups across different cell types, we assigned each cCRE, based on the target signal matrix, to one of the reference clusters. To achieve that, the target PCA signal matrix was first normalized to the reference PCA signal matrix using quantile normalization (initial QTnorm) to mitigate complex technical biases that could result in incorrect cCRE assignments. The Euclidean distance between each cCRE’s target signal vector and the mean signal vector of every reference cluster was then calculated in the PCA space. Each cCRE was assigned to the reference cluster with the smallest Euclidean distance. It is important to note that the initial QTnorm step might introduce noise signals at different PCA space in the data. However, considering that different principal components (PCs) are independent, and the noise signal is expected to randomly appear across different PCs for each cCRE, we reasoned that the majority of PCs would still contain accurate signals, enabling correct cCRE assignment.

The third step of JMnorm involves normalizing the target signal matrix against the reference signal matrix within each cluster in the PCA space. This was achieved by using QTnorm to normalize each principal component separately, removing both simple and complex technical biases that may be present in the target signal matrix. Here, we assumed that cCREs within the same cluster shared the same epigenetic state in both reference and target, and thus had the same signal distributions. However, since the number of cCREs within each cluster could be different between reference and target, they might still have different global distributions. The cCRE clustering followed by within-cluster QTnorm is one of the key distinctions between JMnorm and traditional QTnorm, effectively addressing the major limitation of QTnorm, which forces all cell types to have identical global signal distributions after normalization.

In the fourth step, we reconstructed the normalized target signal matrix in the original signal space. To accomplish that, a dot product was performed between the normalized target PCA signal matrix and the transposed PCA rotation matrix that was learned in the first step.

### Quantification of the similarity of cross-feature correlations

We assessed the ability of various methods to preserve and transfer cross-feature correlation information from reference signal matrices to post-normalization signal matrices of target cell types. Specifically, we calculated the cross-feature correlation matrix for each post-normalization signal matrix of the target cell types and the original reference signal matrix. We then computed the mean squared error (MSE) between each target cell type’s correlation matrix and the reference correlation matrix. Lower MSE values indicate the normalization method can better preserve and transfer cross-feature correlation information from the reference signal matrix to the post-normalization signal matrices of the target cell types. For the cross-species normalization comparison (Figure 2F), S3norm’s cross-feature mode was employed for S3norm normalization that only uses the peak regions and background regions from the reference and target datasets in two species to learn the transformation model. Conversely, MAnorm was excluded because it requires the common peak regions between reference and target datasets to learn its transformation model, which was not available for cross-species normalization.

**Figure 2.**
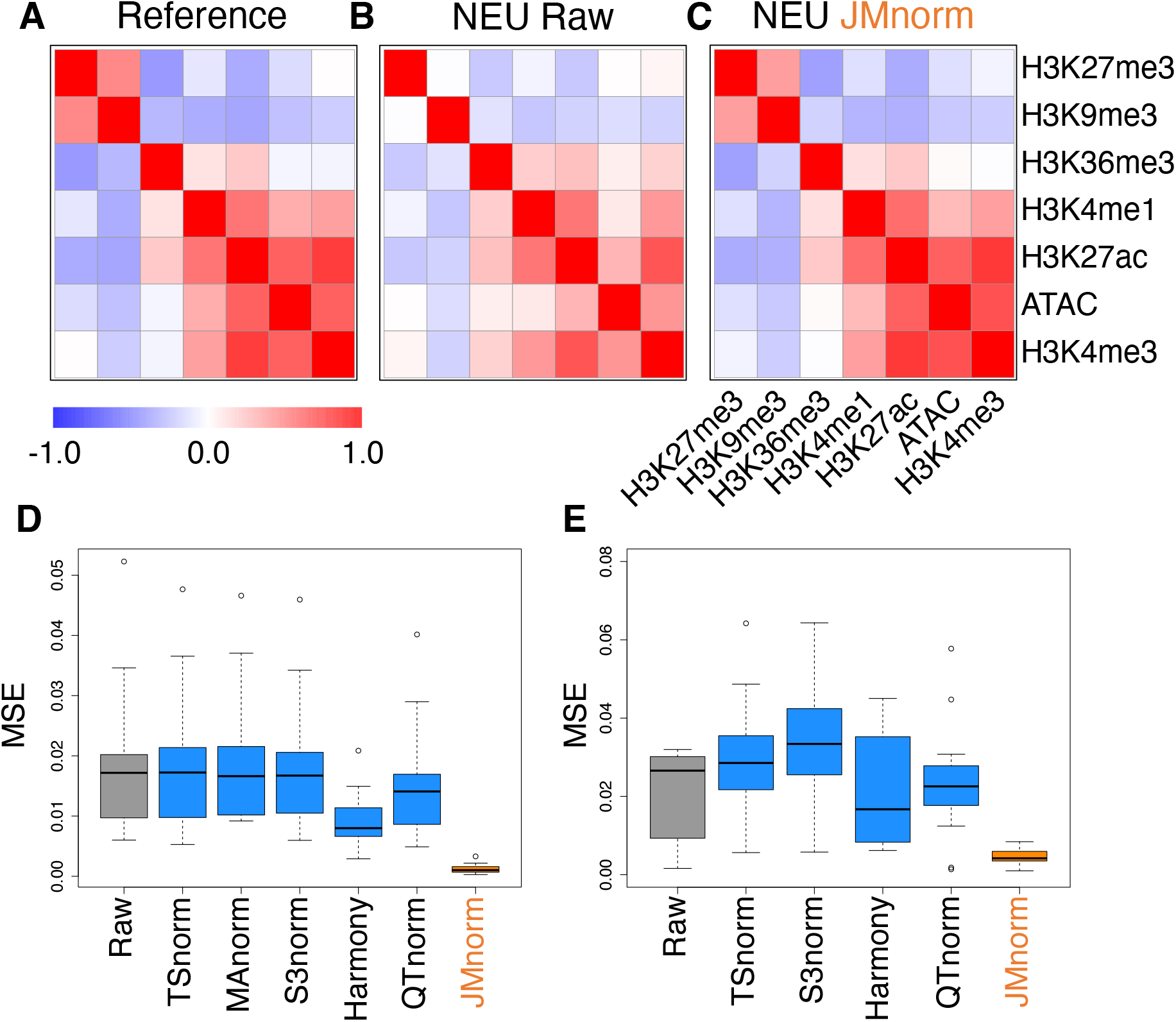
Evaluation of cross-feature correlation preservation. (A) Reference signal cross-feature correlation matrix. (B) Raw signal cross-feature correlation matrix in target neutrophil (NEU) cell type. (C) JMnorm-normalized signal cross-feature correlation matrix in target NEU cell type. (D) A boxplot of MSEs between human target and reference correlation matrices for the six normalization methods. (E). A boxplot of MSEs between correlation matrices of mouse target cell types and the human reference for the five normalization methods.

### Quantification of the preservation of combinatorial patterns across different cell types

To evaluate the preservation of combinatorial patterns for different epigenetic features, we first combined the signal matrices of different cell types and applied a K-means clustering method to group the data from different cell types into distinct clusters. To ensure the robustness of this analysis, we used various numbers of clusters (K = 20, 30, 40, 50) in the K-means clustering step. We then quantified the mixing of various cell types in the clustering results using the average silhouette width (ASW), a metric ranging from 0 to 1 (44). Lower ASW values indicate a better mixing of data points across different cell types within each cluster relative to the between cluster distance.

To measure the proportion of combinatorial patterns that are robustly identified across all cell types, we first performed K-means clustering (K = 30) for each cell type independently. Next, we generated the mean signal vector for each cluster in every cell type and combined all mean signal vectors into a single matrix. We then clustered the mean signal vectors using a second round of K-means clustering (K = 30) (Figure 3D). The output clusters containing at least one mean signal vector from all cell types were defined as robust clusters. We used the proportion of the robust clusters to evaluate different normalization methods. A higher proportion of robust clusters indicates that the normalization method is more effective in generating consistent combinatorial patterns across different cell types. We repeated this process multiple times with different random seeds to ensure the robustness of our conclusions.

**Figure 3.**
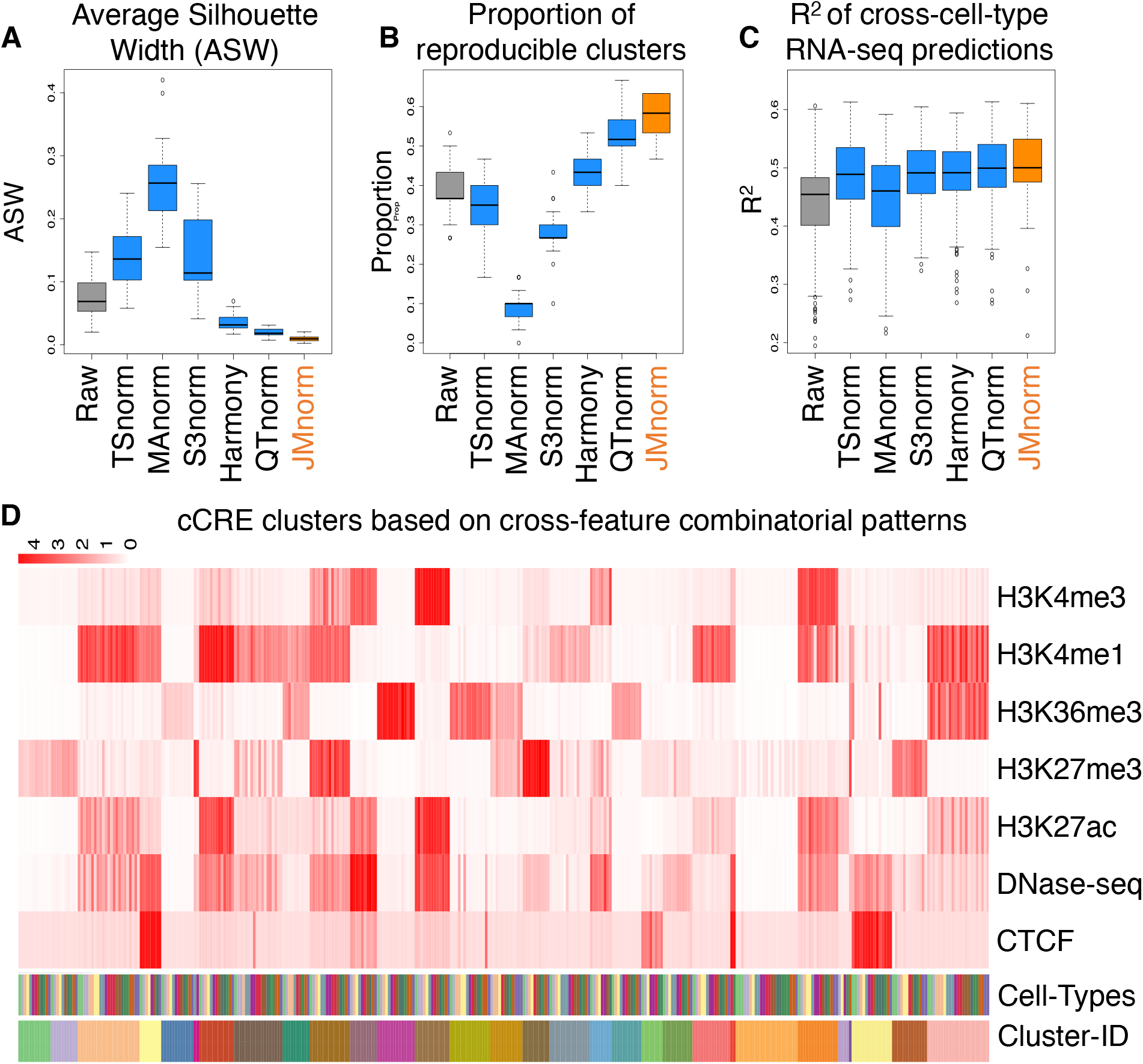
Evaluation of consistency of cross-feature combinatorial patterns and gene expression predictions across cell types. (A) A boxplot of Average Silhouette Widths (ASWs) for the six normalization methods demonstrating quality of mixing of cCREs across different cell types in clustering outputs (K-means: K=30). (B) A boxplot of proportions of cCRE clusters that are reproducible in all cell types for the six normalization methods. (C) A boxplot of R² between observed and predicted RNA-seq values expressed as log2 transcripts per million (log2TPM) for the six normalization methods. Linear regression model with gene grouping was used for the predictions. (D) Top. A heatmap of the average signals at cCRE clusters for seven epigenetic features in 24 EpiMAP cell type groups. Clusters are ordered by the K-means cluster label (K = 30). Bottom. The heatmaps with distinct colors represent labels for different cell types and K-means clusters.

### Cross-cell type RNA-seq predictions

Integration of various epigenetic features, such as DNA accessibility and histone modifications, has been demonstrated to be an effective approach for prediction of gene expression levels. Since RNA-seq data is independent from epigenomic data, we hypothesized that properly normalized epigenetic signal matrices could better preserve the combinatorial patterns of epigenetic events across different cell types, thus enhancing the transferability of RNA-seq prediction models based on these combinatorial patterns and improving the accuracy of gene expression predictions across different cell types. Following the model design proposed by Xiang et al. (3), we used the average signals of eight epigenetic features at both proximal regions (TSS +/− 1kb) and distal cCRE regions (TSS +/− 500kb, excluding proximal regions) for all genes as predictors to train regression models for RNA-seq log2 transcripts per million (log2TPM) predictions. The RNA-seq signals for different cell types were normalized by QTnorm against the average RNA-seq log2TPM across all cell types. To ensure the robustness of our evaluation, we tested three different models: a linear regression model (LM), a gradient boosting regression model (GBM) (45), and a linear regression model with gene grouping (LM-gene-grouping) (3). For gene grouping, we divided the gene set into four groups based on their average expression levels and standard deviations: (1) consistently low (mean < 0.5, sd < 0.2), (2) differentially low (mean < 0.5, sd >= 0.2), (3) differentially high (mean >= 0.5, sd >= 0.2), and (4) consistently high expression (mean >= 0.5, sd < 0.2) across cell types. For each evaluation run, we randomly selected 80% of protein-coding genes in one training cell type to train the model, then used the remaining 20% of protein-coding genes in another testing cell type for model evaluation. Model performance was evaluated using R^2^, with higher R^2^ values indicating more accurate cross-cell type RNA-seq predictions. We repeated this comparison multiple times with different random seeds to ensure the robustness of our conclusions.

### Comparison of the post-normalization signal consistency between biological replicates

We reasoned that improved normalized signals would result in increased consistency between biological replicates. Therefore, we first independently normalized the signals of different replicates. Then, we employed two metrics to quantify signal consistency: (1) the R² values, which were computed between the post-normalization signal vectors of each pair of biological replicates, and (2) the Jaccard index values, which were calculated between the peak-calling results derived from the post-normalization signals of different replicates. Higher values for both R² and Jaccard indexes indicate better post-normalization signal consistency.

### Peak calling from cCRE signal matrices

To compare normalization methods using peak calling results for CTCF ChIP-seq and DNase-seq, we employed an iterative Z-score-based approach, akin to the hotspot peak calling method (46, 47), to call peaks from post-normalization cCRE signal matrices. For a specific epigenetic feature in a given cell type, we first converted normalized signals in all cCREs into Z-scores. We then selected cCREs with false discovery rate (FDR) adjusted p-values >= 0.1 to establish a second-round background model. Next, we recalculated the Z-scores and corresponding p-values for all cCREs using this updated background model. The cCREs with FDR adjusted p-values < 0.1 were used as the output peak list for that chromatin feature in each cell type, which was ultimately utilized in downstream evaluations.

Given that the number of peaks can vary significantly depending on statistical thresholds and methods, we applied a non-parametric, rank-based method to identify the same number of CTCF peaks for each cell type across different normalization methods. Specifically, we first used the iterative Z-score-based strategy to call CTCF peaks for a specific cell type across different normalization methods. The number of peaks (N_max_) for a specific cell type was determined based on the maximal peak count observed across the methods. Then, for different normalization methods, we used the top N_max_ cCREs based on the normalized CTCF signals as the CTCF peaks for the specific cell type.

### Differential peak calling from cCRE signal matrices

As described in the “Peak calling from cCRE signal matrices” section, for different normalization methods, we applied the rank-based method to identify the same number of differential peaks for DNase-seq datasets in different cell-types and glucocorticoid receptor (GR, gene symbol NR3C1) ChIP-seq datasets in A549 cell type with and without dexamethasone (Dex) treatment. Specifically, the top N peaks based on log2 fold change of DNase-seq signals in a pair of cell-types or an absolute value of log2 fold change between Dex-treated and untreated control conditions were defined as differential peaks and used for the downstream differential peak analysis.

### Comparison between CTCF peaks and orthogonal data

To evaluate the ability of normalized signals to accurately capture true biological events, we compared CTCF peak sets derived from different post-normalization CTCF signal vectors with orthogonal data sets. For different normalization methods, the same number of CTCF peaks was generated using the iterative Z-score-based peak calling followed by a rank-based method, as detailed in the “Peak calling from cCRE signal matrices” section. We then evaluated CTCF peaks using three types of orthogonal data: (1) YY1 (CTCF cofactor) peak (48), (2) TAD boundaries (49–51), and (3) the CTCF binding site motifs (52, 53).

We first compared the enrichment of CTCF peaks in YY1 peak regions. We reasoned that true CTCF peaks should exhibit greater enrichment in functionally related regions. Specifically, we divided the CTCF peaks into three distinct groups: (1) CTCF peaks shared by QTnorm / Harmony and JMnorm methods, (2) CTCF peaks uniquely called from JMnorm data, (3) CTCF peaks uniquely called from QTnorm / Harmony data. The enrichments were calculated as follows:

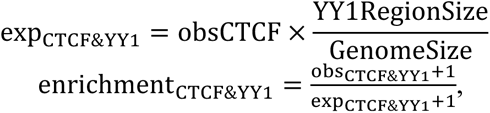

where (obsCTCF) represents the number of CTCF peaks in each group, (YY1RegionSize) represents the total number of base-pairs in the genome covered by YY1 peaks, (GenomeSize) represents the hg38 genome size, (obs_CTCF&YY1_) represents the observed number of CTCF peaks that intersect with YY1 peak regions by at least one base-pair. Due to the limited availability of YY1 peak datasets in some cell types, we pooled available YY1 peaks creating a single unified YY1 peak set for the YY1-CTCF peak intersection enrichment comparisons. Similarly, due to the availability issue for TAD boundary sets, we employed a unified TAD boundary region set for TAD boundary-CTCF peak intersection enrichment comparisons (Supplementary Table 1).

We then compared the proportion of CTCF peaks containing CTCF motif (Jaspar ID: MA0139.1) for the data normalized by different normalization methods. We used FIMO (54) to assess whether CTCF peak contained CTCF motif using a q-value threshold of <= 1e-03 for motif identification. The CTCF peak set was partitioned into the same three groups as described earlier for the CTCF-YY1 enrichment comparison.

### Clustering cCRE based on cross-cell type DNase-seq patterns

We employed the Snapshot package with default settings to cluster cCREs based on DNase-seq signal patterns across different cell types. Snapshot was chosen for its ability to efficiently identify smaller clusters, automatically determine the ideal cluster number, and consider signal correlations across cell types when grouping cCREs into separate clusters using an indexing strategy (8). To quantify the signal-to-noise ratios for the Snapshot output clusters, we defined “Signal” as the top quantile (100%, 90%, 80%, 70%, 60%) signal and “Noise” as the bottom quantile (10%, 20%, 30%, 40%, 50%) signal in the meta-cluster heatmap. We calculated the signal-to-noise ratio using different combinations of “Signal” and “Noise” and summarized the results in Figure 6E. Statistical significance of the differences was calculated using the Paired Wilcoxon test.

### Identification of cell-type-specific DHSs

For each cell type within a cell type pair, we initially identified 10,000 cell-type-specific DHSs from the post-normalization DNase-seq signal vectors using different normalization methods, following the strategy detailed in the “Differential peak calling from cCRE signal matrices” section. Then, using the *bedtools intersect* function with default parameters (55), these cell-type-specific DHSs were categorized into three groups: JMnorm-uniquely identified, QTnorm-uniquely identified, and those shared by both JMnorm and QTnorm methods.

### Identification of differential glucocorticoid receptor (GR) ChIP-seq peaks after dexamethasone (Dex) treatment

For GR motif analyses and comparison of human phenotype term enrichments, we used the top 2,000 differential peaks, identified based on the absolute value of the log2 fold changes between GR signals in A549 cells with and without Dex treatment. For different normalization methods, we analyzed the same number of differential GR ChIP-seq peaks. We computed GR motif (JASPAR ID: MA0113.1) scores in differential peaks as the −log10 p-value using the FIMO (54) motif scanning algorithm. The method for calculating the human phenotype term enrichment score is described in the “Human Phenotype term enrichment by GREAT analysis” section.

### Human Phenotype term enrichment by GREAT analysis

To quantify enrichment of human phenotype terms for genes associated with distinct peak sets including cell-type-specific DHS and differential GR peaks post-Dex treatment, we utilized the rGREAT package (56) (57). In this study, proximal regions were defined as TSS −5 kb to +1 kb, and distal regions were defined as proximal regions ± 100 kb. To focus on more specific terms, we excluded human phenotype terms linked with more than 1,000 genes to eliminate overly general associations.

To compare the enrichment of human phenotype terms in cell-type-specific DHSs or differential GR ChIP-seq peaks after Dex treatment, we assumed that DHSs that are shared across normalization methods or differential GR ChIP-seq peaks that are identified using TSnorm_cbg corresponded to epigenetic signals representing true biological events. We identified the top 30 enriched human phenotype terms for these shared DHSs or differential GR ChIP-seq peaks. Next, for each normalization method, we quantified enrichments of the top 30 terms in cell-type-specific DHSs uniquely identified by each method and used these enrichments as a metric for method comparisons. Since relatively few differential GR peaks were unique to different normalization methods impairing the reliability of significance calculation of the human phenotype term enrichments, we compared performance of the normalization methods using enrichment of the top 30 terms in all differential GR peaks identified by each method instead.

### Using Harmony for cross-cell type multi-feature batch corrections

The Harmony method (58), originally designed for single-cell batch correction, transforms the input signal matrix to PCA space for batch correction. During this process, it modifies the signal matrices of both reference and target cell types in each run. As a result, in order to correct technical biases for signal matrices across cell types against a single reference matrix, both reference matrix and matrices for all cell types would need to be included in a single Harmony run, which would consume an excessive amount of computational resources. To address this issue, for each Harmony run, we assigned five times greater weight to the reference signal matrix than to the signal matrix of an individual target cell type. We reasoned that potential technical signal biases of the reference signal matrix would outweigh the biases of individual target cell type. This approach allowed for batch correction of data across all cell types against the same reference signal matrix without requiring the inclusion of all cell types’ data in a single Harmony run.

## RESULTS

### JMnorm overview

The goal of JMnorm is to simultaneously normalize multiple epigenetic features across cell types, species, or experimental conditions by leveraging information from functionally correlated features. The input for JMnorm is signal matrices comprising data for multiple epigenetic features for a desirable number of regulatory regions in two or more cell types or experimental conditions (reference and target cell types or conditions). To develop the method, we utilized human and mouse ENCODE cCREs (2), comprehensive collections of genome-wide regulatory elements generated in multiple cell types in the two species.

The method consists of four key steps (Figure 1). In step 1, orthogonal transformation, JMnorm converts the correlated components of multi-dimensional epigenetic signal matrices for both reference and target cCREs into mutually independent PCA dimensions. This process generates corresponding PCA matrices, effectively preserving the relationships between functionally related epigenetic features (Figure 1A) (59). In the subsequent steps 2 and 3, JMnorm performs data normalization within the PCA space, which simultaneously normalizes all features and transfers the cross-feature patterns from the reference signal matrix to the post-normalization signal matrices of the target cell types or conditions. We anticipate that this strategy can improve the performance of downstream prediction models that utilize the cross-feature patterns for tasks such as gene expression prediction (60–62), enhancer-promoter interaction prediction (63–65), or epigenomic data imputation (66–70), especially for cross-cell-type predictions. Specifically, in step 2, cCRE clustering, JMnorm first clusters (N) cCREs based on the reference PCA matrix into (M) reference clusters (M << N). It then assigns each of the individual target cCREs into the nearest reference cluster based on signal similarity in the PCA space (Figure 1B). In step 3, within-cluster normalization, JMnorm normalizes the target cCRE signals (PCA-transformed) to the corresponding reference cCRE signals (PCA-transformed) using quantile normalization within each cluster (Figure 1C). When performing cCRE clustering (step 2) followed by within-cluster QTnorm (step 3), we assume that (1) the combinatorial patterns of epigenetic features are conserved across cell types or conditions and (2) within each cCRE cluster, the signal distributions of PCA matrices are also conserved across cell types or conditions. It is important to note that the number of target cCREs assigned to each reference cluster can vary for different target datasets, thereby resulting in different global signal distributions across cell types or conditions after normalization. Therefore, this strategy can circumvent potential biases inherent in QTnorm, which forces identical global signal distributions across diverse datasets. Once steps 1-3 are completed, JMnorm transforms the normalized target signal matrix from the PCA space back to the original signal space in step 4 (Figure 1D). The details of these steps can be found in the Materials and Methods section.

### Evaluation of preservation of cross-feature correlation

We evaluated the performance of JMnorm relative to a panel of other normalization methods (TSnorm, MAnorm, S3norm, QTnorm), which are frequently applied as initial data normalization strategies for various downstream analyses. We also included Harmony, a widely used batch correction method for single-cell data. Similar to JMnorm, Harmony utilizes PCA transformation to harness information from correlated features within multi-dimensional data (58).

The first evaluation was to compare the performance of JMnorm and other normalization methods for their ability to preserve the cross-feature correlation among different epigenetic features across multiple cell types. While it is possible to use data from one particular cell type as a reference for normalizing data from other cell types, this approach could introduce biases from the data of the chosen cell type. To circumvent such potential biases, we generated a reference dataset by averaging each epigenetic feature across all cell types. Next, we used the selected methods to normalize raw signal matrices of target cell types against the reference signal matrix. The resulting post-normalization signal matrices were then used to compute the cross-feature correlation matrices.

We first inspected the cross-feature correlation matrices derived from the reference, the raw data, and JMnorm-normalized data from neutrophil (NEU) cells, respectively (Figure 2A, B, and C). We observed that the JMnorm-derived correlation matrix was more similar to the reference correlation matrix than to the correlation matrix of the raw data. Specifically, correlation matrices derived from both JMnorm and reference signal matrices exhibited strong positive correlations between features within the active chromatin feature group (H3K27ac, H3K4me3, H3K4me1, ATAC-seq) and the repressed chromatin feature group (H3K27me3 and H3K9me3) (Figure 2A and C). Conversely, we observed a substantial negative correlation between features in active and repressed chromatin groups (Figure 2A and C) (1, 2). In contrast, the aforementioned correlation relationships are much weaker in the raw data (Figure 2B), likely due to the technical biases in the raw data. These results suggest that JMnorm is effective in reducing technical biases and preserving and transferring the cross-feature correlation information, which is better aligned with our prior knowledge, from the reference to the target cell type.

We then compared the panel of methods by measuring the MSEs between cross-feature correlation matrices of the normalized data and the reference across different cell types. JMnorm-derived correlation matrices better preserved the information in the reference correlation matrix than those derived from other methods, as indicated by its significantly lower MSEs (Paired Wilcoxon test between JMnorm and the second-best performing method, p-value = 3.81e-06; Figure 2D).

Furthermore, the superior performance of JMnorm also holds true in cross-species normalizations, when mouse cCRE signal matrices of different cell types were normalized against the human reference signal matrix. Specifically, JMnorm produced significantly lower MSEs than other methods (Paired Wilcoxon test between JMnorm and the second-best performing method, p-value = 2.13e-04) when comparing the correlation matrices of each mouse target cell type to the human reference (Figure 2E).

Since experimental data for large panels of epigenetic features might not always be available, we also assessed JMnorm’s performance with fewer features. As demonstrated in Supplementary Figure 1, JMnorm can effectively preserve and transfer cross-feature correlation information when normalizing datasets containing as few as two epigenetic features.

### Evaluation of consistency of cross-feature combinatorial patterns across cell types

Combinatorial patterns of epigenetic features, called epigenetic states, reflect functionally relevant interactions between DNA accessibility, histone modifications, and transcription factor binding under specific cellular and experimental conditions. They are often used to accurately infer transcriptional states and interpret the function of non-coding genetic variants (2, 23–26, 60–62). Previous studies have shown that epigenetic states are maintained across different cell types or even species (4). With the expanding variety of epigenetic features and functional element annotations along with the growth in profiled cell types, there is a growing need for normalization methods that can effectively integrate and preserve recurring combinatorial patterns within the epigenetic states across diverse cell types.

We first compared the panel of methods for their ability to harmonize signal matrices of various epigenetic features across multiple cell types. A better normalization method should more consistently preserve common combinatorial patterns in different cell types and thus allow better data integration across cell types. To quantify the degree of preservation of common combinatorial patterns, we first pooled cCRE signal matrices of seven epigenetic features from each pair of different cell types and identified the combinatorial patterns by clustering cCREs using K-means (K = 30, 20, 40, 50). We then measured the ASW score, which ranges from 0 to 1 and a lower score indicates better sharing of combinatorial patterns between cell types (44). As shown in Figure 3A and Supplementary Figure 2, JMnorm has significantly lower ASW scores than other methods (Paired Wilcoxon test between JMnorm and the second-best performing method, p-value = 3.05e-05, 3.05e-04, 2.14e-04, 6.23e-03 for different K values in K-means clustering), indicating that the multi-feature signal matrices normalized by JMnorm achieved better integration among cell types within each cCRE clusters.

Next, we clustered all cCREs from all 24 cell types groups available in EpiMAP database and used the resulting cCRE clusters (Figure 3D) to compare different normalization methods, using the proportion of robust cCRE clusters (4) as a metric of global preservation of epigenetic feature patterns across different cell types. Here, a robust cCRE cluster is defined as the cCRE cluster displaying high consensus across all cell types. A higher proportion of robust cCRE clusters would indicate a better preservation of combinatorial feature patterns across cell types (see method section for details). JMnorm-normalized signal matrices resulted in a significantly higher proportion of robust cCRE clusters (Paired Wilcoxon test between JMnorm and the second-best performing method, p-value = 2.60e-04) than those obtained by other methods (Figure 3B). Results of these comparisons demonstrate that JMnorm outperforms other methods in integrating multi-feature signal matrices and preserving cross-feature combinatorial patterns across different cell types.

### Evaluation of cross-cell-type gene expression predictions

Since JMnorm integrates the multi-feature signal matrices and preserves cross-feature combinatorial patterns across different cell types better than other methods, we anticipated that the gene expression prediction models utilizing data normalized by JMnorm would also exhibit improved performance in cross-cell-type predictions. Therefore, we compared normalization methods by their ability to improve cross-cell type gene expression predictions. We trained three types of regression models to learn the quantitative relationships between seven epigenetic features and RNA-seq data: a linear regression model (LM), a gradient boosting regression model (GBM) (45), and a linear regression model with gene-grouping that is based on cross-cell type average expression levels and standard deviations (LM-gene-grouping) (3). The details about training and testing of these regression models are described in the Materials and Methods section. Briefly, we randomly divided protein-coding genes into two groups: 80% of genes were used for model training (Training-Genes) and 20% of genes were used for model evaluation (Testing-Genes). The regression models were trained using the signals of Training-Genes in one Training-Cell-Type and evaluated by R-squared (R^2^) using the signals of Testing-Genes in another Testing-Cell-Type, representing the most challenging cross-cell type hold-out gene prediction setting in gene expression prediction tasks. As shown in Figure 3C and Supplementary Figure 3, JMnorm-normalized data had significantly higher R^2^ than others normalization methods across all three regression methods and different cell-type-pairs (Paired Wilcoxon test between JMnorm and the second-best performing method, p-values are LM: 6.16e-06, GBM: 1.48e-08, LM-gene-grouping: 2.54e-07). These results demonstrate that JMnorm-normalized epigenetic signals improve performance of cross-cell type gene expression prediction models, suggesting that the JMnorm-normalized epigenetic data are more consistent with gene expression levels, an orthogonal biological data type, than those normalized by other methods.

### Evaluation of signal consistency between biological replicates

We next evaluated different methods based on consistency of post-normalization signal strengths between biological replicates, measured by R^2^. A better normalization method should result in higher signal consistency between the independently normalized replicates. We examined the replicate consistency of the post-normalized H3K27ac ChIP-seq signals across 9 different cell types. Six other features were used for JMnorm normalization of H3K27ac ChIP-seq data. JM norm-normalized data had significantly higher R^2^ values than data normalized by other methods (Paired Wilcoxon test between JMnorm and the second-best performing method, p-value = 1.95e-03) (Figure 4A). Moreover, JMnorm-normalized data had the highest or the second highest R^2^ values across all 7 examined epigenetic features (Figure 4A and Supplementary Figure 4), especially those that are more likely to reflect cell-type-specific epigenetic events such as ATAC-seq, H3K27ac, and H3K4me1 (1).

**Figure 4.**
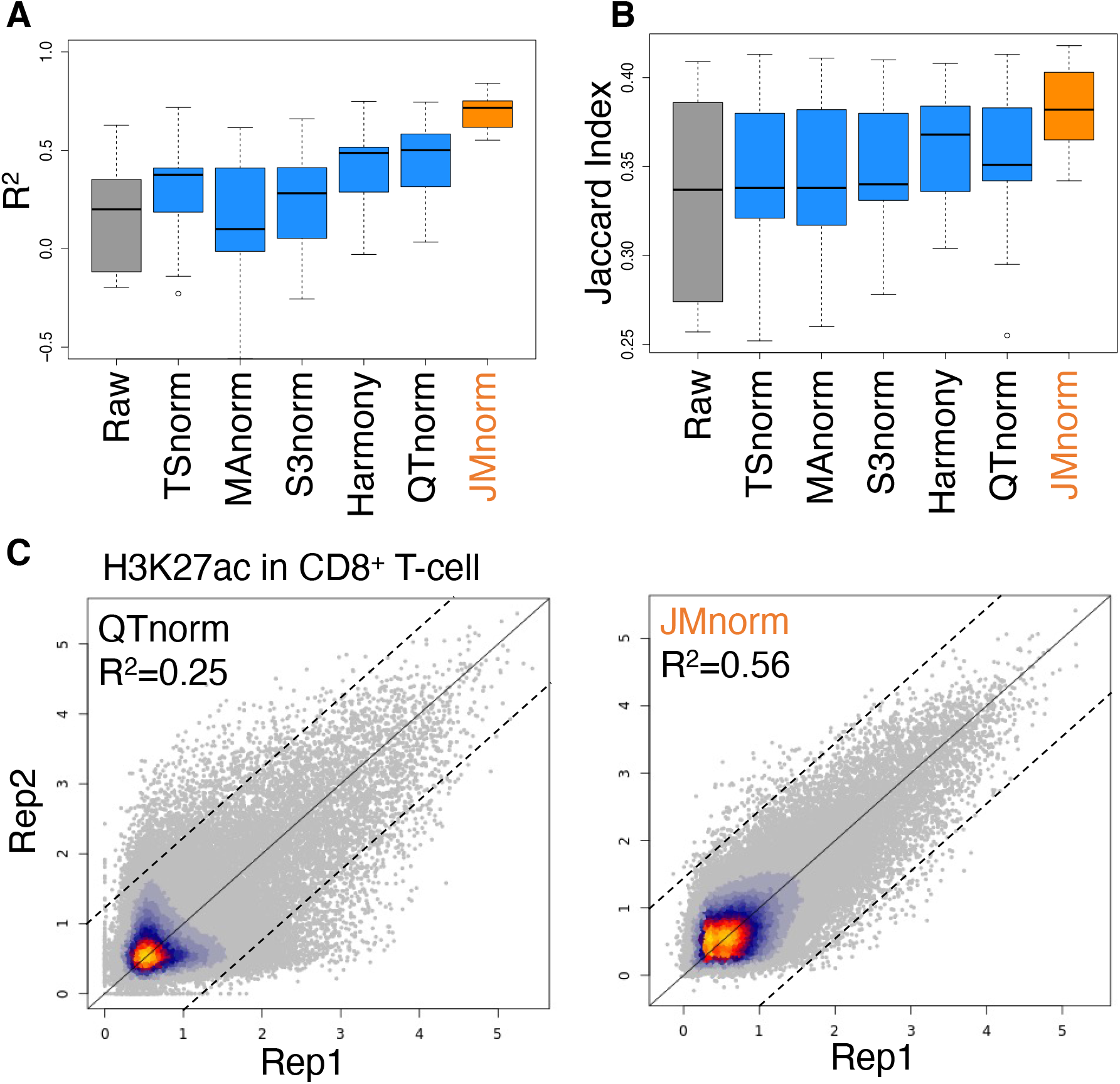
Evaluation of the post-normalization signal consistency between biological replicates. (A) A boxplot of R^2^ values between biological replicates in multiple cell types for the six normalization methods. (B) A boxplot of Jaccard indexes comparing peak calling results between biological replicates of multiple cell types for the six normalization methods. (C) Scatterplots of H3K27ac ChIP-seq signal in CD8^+^ T-cells in two biological replicates normalized by QTnorm (left) and JMnorm (right). Data are shown on log2 scale. Bright orange and gray colors indicate higher and lower data point density, respectively.

As an additional metric, we assessed the replicate consistency of enriched peak calling results for H3K27ac across 9 cell types. To this end, the same peak-calling method was applied to post-normalized signals generated by the panel of normalization methods, and an equal number of called peaks was identified for evaluation of replicate consistency using the Jaccard index. Besides the signal strengths consistency, JMnorm’s peak calling results also had higher replicate consistency as indicated by significantly higher Jaccard index values compared to peaks generated from data normalized by other methods (Paired Wilcoxon test between JMnorm and the second-best performing method, p-value = 4.5e-03; Figure 4B).

We then generated scatterplots of post-normalization signals at individual cCREs for the two biological replicates from the H3K27ac ChIP-seq experiment in CD8^+^ T-cells (Figure 4C and Supplementary Figure 5). Compared to other methods, the scatterplot for JMnorm showed considerably less deviation from the diagonal line (indicating a higher R^2^), especially for the cCREs with lower or only noise-like signals indicated by the red arrow at bottom left corner of the scatterplots.

These results suggest that by leveraging information from functionally correlated features, JMnorm can more effectively improve replicates consistency by reducing technical noise and preserving true epigenetic signals than other normalization methods.

It is important to note that JMnorm performance is not universally superior to all other methods for normalization of all examined epigenetic features. For example, since H3K4me3 is a canonical marker of gene promoters, its ChIP-seq signals have similar global distributions across different cell types. This type of signal distribution is expected to be suitable for QTnorm, which enforces identical post-normalization signal distributions. Hence, QTnorm exhibited a slightly higher R^2^ between biological replicates for H3K4me3 ChIP-seq signal than JMnorm (Supplementary Figure 4). Based on the overall performance, we conclude that JMnorm surpasses other normalization methods in enhancing signal and minimizing technical noise and improves consistency between biological replicates.

### Evaluation of quality of peak calling results

Based on the performance evaluations results above, JMnorm, QTnorm, and Harmony outperformed the other three methods. This outperformance is expected since JMnorm incorporates the strengths of both QTnorm and Harmony: as QTnorm, it minimizes the complex technical biases by equalizing the signals of the same rank, and as Harmony, it combines the highly correlated variables in the PCA space. To more closely evaluate the six normalization methods, we compared their performance by the quality of peak calling results (Figure 5 and Supplementary Figure 6 and 7). We hypothesized that better normalization could improve the accuracy of peak calling results by reducing both false positives and negatives, thereby yielding peaks more closely associated with true biologically relevant events.

**Figure 5.**
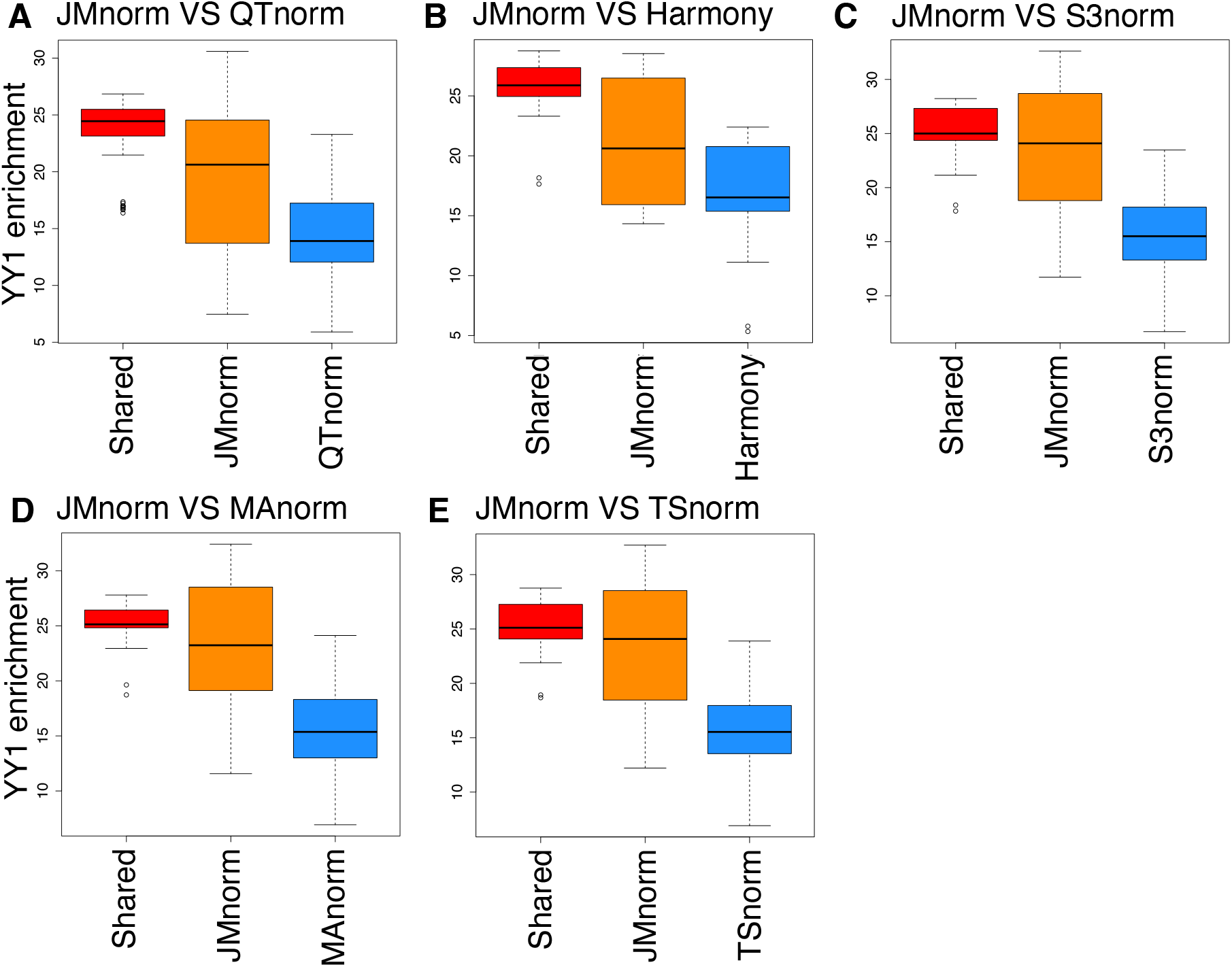
Evaluation of quality of TF peak calling results. Box plots comparing JMnorm’s and five other methods’ performance as determined by CTCF peak enrichment at YY1 peak regions. (A) Comparison between JMnorm and QTnorm, (B) JMnorm and Harmony, (C) JMnorm and S3norm, (D) JMnorm and MAnorm, and (E) JMnorm and TSnorm. Red box plots represent enrichments for CTCF peaks that are shared between JMnorm and a corresponding alternative method. Orange box plots represent enrichments for CTCF peaks uniquely identified by JMnorm. Blue box plots represent enrichments for CTCF peaks uniquely identified by the respective alternative method.

For this test, we first selected TF ChIP-seq data for CTCF (71) and YY1 (48) because the role of these TFs in transcription regulation at gene promoters, enhancers, and topologically associating domain (TAD) boundaries is well understood and their DNA-binding motifs had been extensively characterized. Specifically, CTCF and YY1 often co-occupy the same genomic regions at the promoters and enhancers of active genes (48), whereas TAD boundaries are uniquely bound by CTCF (49–51). Moreover, approximately 80% of previously validated CTCF ChIP-seq peaks contain CTCF binding motifs (52, 53). To evaluate the quality of CTCF ChIP-seq peak calling results, we applied the same peak-calling method to the post-normalized signals generated by the five normalization methods, yielding an equal number of CTCF peaks for each method (see Materials and Methods section for more details). The resulting CTCF peaks were evaluated using the following three metrics: (1) enrichment at YY1 peak regions, (2) enrichment at TAD boundaries, and (3) fraction of the CTCF peaks with CTCF DNA-binding motif. We found that CTCF peaks, uniquely called using JMnorm-normalized data, exhibited significantly higher enrichment at YY1 peak regions than the ones generated by other methods (Paired Wilcoxon test p-values < 0.05) (Figure 5). Similarly, JMnorm-normalized CTCF ChIP-seq peaks had significantly higher (Paired Wilcoxon test p-values < 0.05) or comparable enrichment at TAD boundaries than other methods’ peaks (Supplementary Figure 6). Lastly, there is no significant difference in proportion of CTCF peaks containing CTCF motifs (Jaspar (72) ID: MA0139.1) with the JMnorm-normalized data relative to other methods’ data (Supplementary Figure 7). In sum, these findings suggest that JMnorm exhibits improved performance in terms of quality and biological significance of the resulting TF ChIP-seq peaks.

We next focused on comparing the JMnorm and QTnorm by the quality of the DNase-seq peak calling results across cell types (Supplementary Figure 8). Both methods are designed to minimize technical biases by equalizing the signals of the same rank. However, a critical technical bias of QTnorm is that it imposes an identical signal distribution across cell types, which might lead peak-calling algorithms to identify the same number of peaks for all cell types. We asked if JMnorm could circumvent this technical bias since it applies QTnorm only within each cCRE cluster, which may contain a different number of cCREs per cell type, thereby leading to different global signal distributions across cell types and varying numbers of peaks for the normalized data.

To this end, we first compared the performance of QTnorm and JMnorm on the same set of DNase-seq data generated in 24 cell types and found that QTnorm calls the same number of peaks across cell types, whereas JMnorm effectively identifies a different number of peaks per cell type (Supplementary Figure 8A). As expected, most cell types close to stem cells exhibited a larger number of peaks (denoted by orange coloring) than other cell types, consistent with prior observations by Meuleman et al. (40) (Supplementary Figure 8B).

We then evaluated the noise level for the two methods. Since QTnorm imposes an identical signal distribution across various cell types, we expected more false positive peaks in the cell types with fewer true biological peaks. Clustering of cCREs based on these signals would reveal noisier cross-cell type patterns with increased presence of weaker signals in many cell types. To test the validity of these expectations, we utilized the Snapshot clustering algorithm to group cCREs using normalized DNase-seq signals (8) and then evaluated the quality of the resulting clusters. As expected, QTnorm clusters exhibited relatively noisier cross-cell type patterns (Supplementary Figure 8C and D) with significantly lower signal-to-noise ratios than JMnorm’s clusters (paired Wilcoxon test p-value = 2.38e-07) (Supplementary Figure 8E) (see Materials and Methods section for more details), indicating that JMnorm-normalized DNase-seq signals had less noise.

Lastly, to determine whether the JMnorm-normalized DNase-seq peaks contained true biologically meaningful information, we conducted pairwise comparisons of post-normalized DNase-seq peaks in different cell types and assessed the enrichment of human phenotype terms (73, 74) relevant to the respective cell types in the differential peaks. We reasoned that cell-type-specific differential peaks would be more likely to reflect true biological differences between cell types and corresponding top-enriched human phenotype terms could be used to highlight the cell-type-specific functional differences. An initial analysis of the differential peaks identified in heart and lung cell types revealed greater enrichment of heart-relevant terms in heart-specific peaks uniquely identified using JMnorm-normalized DNase-seq data than QTnorm-normalized data (Figure 6A). Next, we conducted the same pairwise DNase-seq peak analysis for 100 randomly selected pairs of cell types. Differential peaks unique to JMnorm had a significantly higher enrichment (paired Wilcoxon test p-value < 0.05) in cell-type-specific functional terms than peaks uniquely identified using other methods (Figure 6B-F), indicating that JMnorm-derived differential peaks may more accurately capture true biological information.

**Figure 6.**
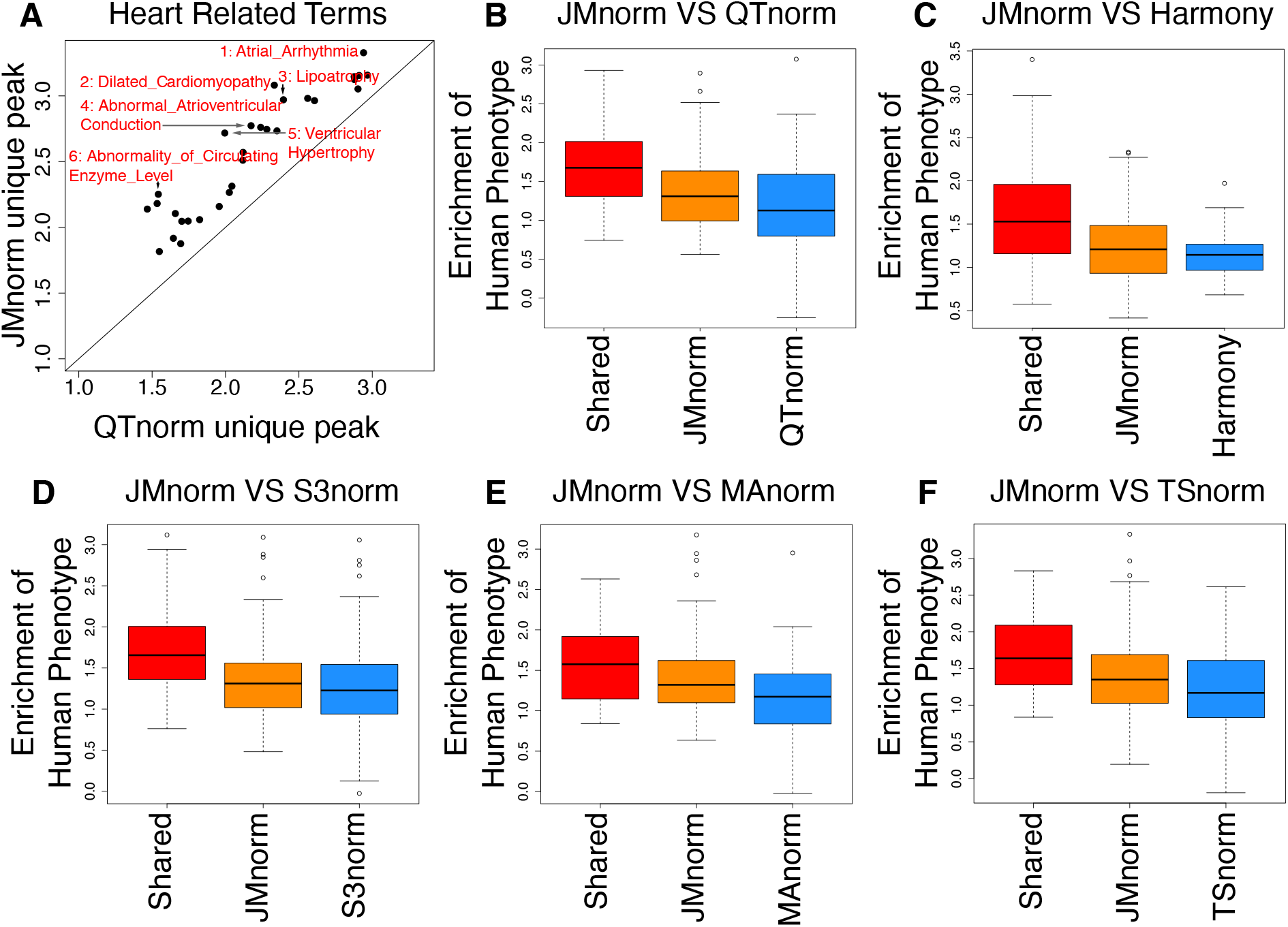
Comparative analysis of JMnorm and other methods using DNase-seq peak calling results. (A) A scatterplot of enrichments of the Human Phenotype terms in heart-specific peaks (relative to lung) uniquely identified by QTnorm (x-axis) or JMnorm (y-axis). The displayed terms are the top 30 most significantly enriched terms identified by both methods. (B) A box plot of Human Phenotype terms enriched in shared and uniquely identified QTnorm’s or JMnorm’s peaks across 100 randomly selected cell type pairs. (C-F) same as (B), for comparison between JMnorm and Harmony (C) / S3norm (D) / MAnorm (E) / TSnorm (F).

These results demonstrate that JMnorm improves the accuracy and biological relevance of the peak calling results.

### Evaluation of quality of differential peak calling in response to perturbations

Small molecule perturbations often make a substantial and global impact on the epigenome (75, 76) presenting a substantial challenge for data normalization. That is because many normalization methods assume that differences in the overall signal (12–14) or common peak regions (16, 17) between different datasets are caused primarily by technical inconsistencies rather than true biological changes in global epigenetic signals. We compared the performance of six normalization methods by the quality of differential glucocorticoid receptor (GR, gene symbol NR3C1) ChIP-seq peak calling results in the context of dexamethasone (Dex) treatment of A549 cells (15) (Figure 7). We selected GR ChIP-seq data for this evaluation because GR is a well-characterized receptor for Dex, GR’s binding to DNA increases globally with transient Dex treatment, and its DNA-binding motif is also well known (78).

**Figure 7.**
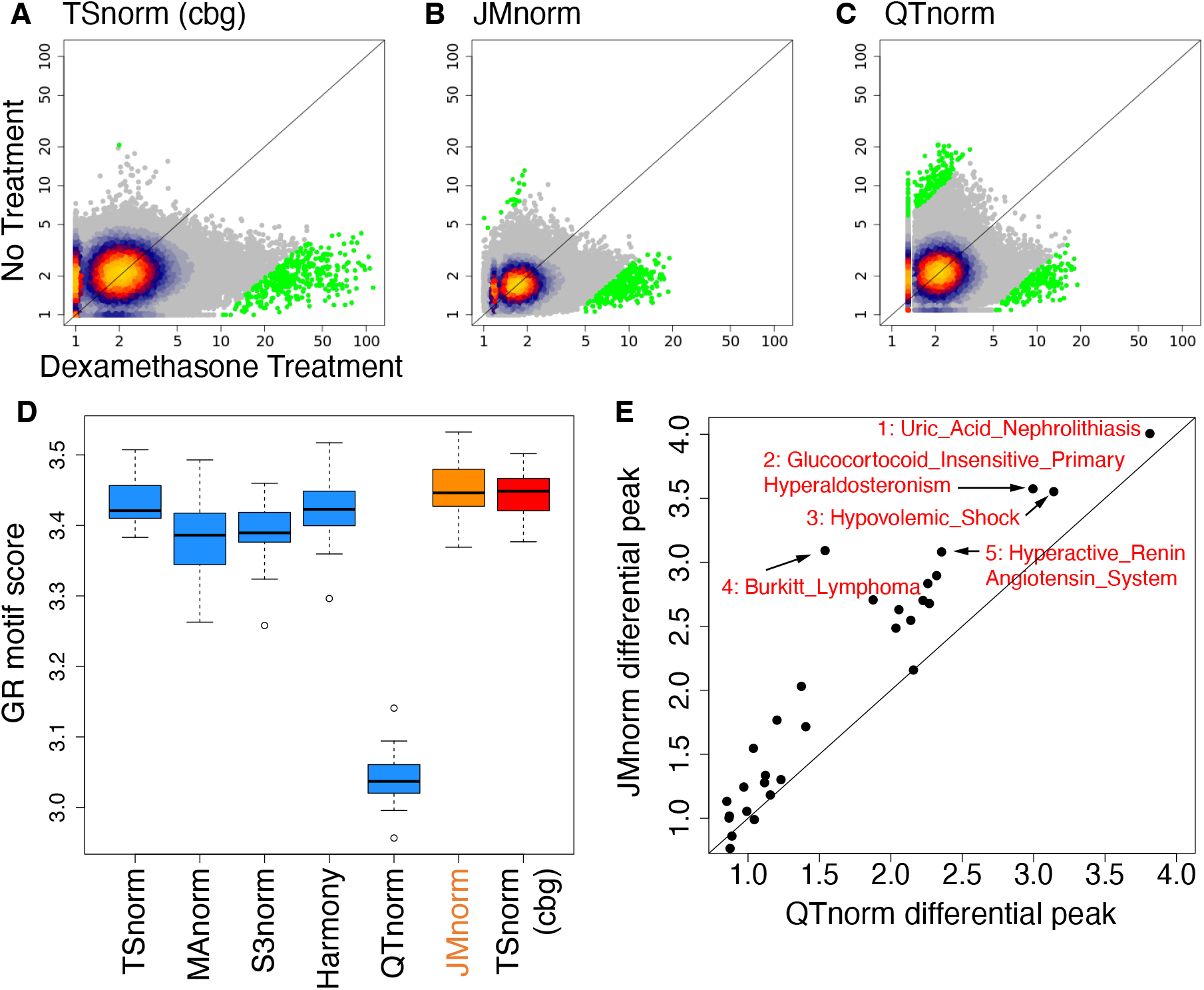
Evaluation of quality of differential peak calling in response to perturbations. (A-C) Scatter plots of GR ChIP-seq signals at ENCODE cCREs with (x-axis) and without (y-axis) Dex treatment for (A) TSnorm (cbg), (B) JMnorm, and (C) QTnorm. Bright orange and gray colors indicate higher and lower data point density, respectively. Green color represents GR ChIP-seq signals for the top 2,000 differential peaks based on absolute value of log2 fold change of signals between Dex treatment and no treatment control. (D) A boxplot of FIMO scores for GR motif (Jaspar ID: MA0113.1) found in differential peaks for the seven normalization methods. (E) A scatterplot of enrichments of the Human Phenotype terms in differential GR ChIP-seq peaks identified by QTnorm (x-axis) or JMnorm (y-axis).

For this analysis, we first normalized the GR ChIP-seq data for both the Dex treatment condition and the no-treatment control using each of the six methods and then identified differential GR peaks using the post-normalization signals. For JMnorm and Harmony, the GR ChIP-seq data were normalized in conjunction with four additional epigenetic features (ATAC, H3K27ac, H3K4me1, and H3K4me3). We benchmarked the performance of different normalization methods relative to TSnorm_cbg. The TSnorm_cbg leverages the same concept as the normalization of ChIP-seq with control (NCIS) (77) method that was specifically developed to address the challenge of normalizing global differences between ChIP and control datasets, by calculating a scale factor based on the information from common background regions between the two. We hypothesized that better normalization could improve the accuracy of differential peak calling results, yielding peaks that are more closely associated with true biologically relevant changes induced by the perturbagen.

As expected, most of the differential GR peaks (top 2,000 differential peaks based on absolute value log2 fold-change between Dex treatment sample and no treatment sample) called using TSnorm_cbg-normalized data were upregulated after Dex treatment (Figure 7A), indicating an increase of GR binding to the genome. For JMnorm, most differential peaks also exhibited increased GR signals (Figure. 7B), similarly supporting our biological understanding. Conversely, only approximately 50% of differential peaks called using QTnorm-normalized data were upregulated with Dex treatment (Figure 7C).

To further characterize the differential GR peaks identified using data normalized by the six methods relative to TSnorm_cbg-normalized data, we evaluated the quality of the GR motif (Jaspar ID: MA0113.1) (Figure 7D) and the extent of Human Phenotype term enrichment (Figure 7E and Supplementary Figure 9) in differential peaks. As expected, GR motif scores and the degree of GR-related process term enrichment were significantly lower in differential peaks called using QTnorm/S3norm/MAnorm-normalized data as compared to those derived from the JMnorm (paired Wilcoxon test p-value < 0.05). These results suggest that an improper matching of overall distribution or signal-to-noise ratio can undermine the association between differential GR peaks and Dex treatment. In conclusion, JMnorm improves the accuracy and biological significance of the differential peak calling results relative to QTnorm/S3norm/MAnorm, and demonstrates a comparable performance relative to the other approaches including the state-of-the-art method (TSnorm_cbg) for normalization of epigenetic datasets with global differences induced by perturbations.

## DISCUSSION

We present a novel approach named JMnorm that simultaneously normalizes multiple epigenetic features across cell types, species, and experimental conditions by leveraging information from functionally correlated features. JMnorm presents several methodological advances over existing normalization approaches that analyze each feature independently and thus may distort relationships between epigenetic features. Specifically, JMnorm normalizes multiple epigenetic features jointly in the PCA space that preserves correlations between different features. Secondly, using a two-step process of initial cCRE clustering followed by within-cluster quantile normalization, JMnorm effectively reduces technical biases without imposing identical global signal distributions across different cell types. Following the principle of Occam’s razor, we have implemented JMnorm using simple yet sufficiently effective statistical techniques. By employing these techniques, we aimed to enhance the robustness of JMnorm and enable its wide application for various downstream analyses. Future investigations could potentially benefit from exploring advanced techniques, such as orthogonal transformation and cCRE clustering. We have demonstrated JMnorm’s improved capabilities by comparing cross-feature correlation matrices before and post-normalization, analyzing the extent of preservation of combinatorial patterns of epigenetic features across different cell types, and evaluating consistency between biological replicates. Furthermore, in several use cases of epigenomic analyses, such as prediction of gene expression, peak calling, and differential TF binding, we showed that JMnorm achieves better results than other methods when being validated against various types of orthogonal biological data. Altogether, these improvements underscore the strength of JMnorm in reducing noise and preserving true biologically meaningful information in epigenomic data.

Given the superior performance of JMnorm across diverse types of epigenetic features and genomic tasks, we anticipate that it can be used across a broad range of applications. In addition to the applications described in this work, a potential application for JMnorm is in normalizing single-cell gene module signal matrices. Single-cell gene expression analyses (80, 81) often involve data transformation that converts relatively noisy individual gene expression information into gene module scores (82–86). By leveraging information from correlated gene modules, JMnorm may provide a more accurate representation of cells or meta cells in different gene module spaces across various cell groups, conditions, or time points.

Because JMnorm normalizes data by leveraging information from functionally correlated datasets, higher correlations among the features can result in better improvements with normalization. To help users more effectively select features with high correlations for JMnorm analyses, we computed pairwise cross-feature correlations for 538 epigenetic features in K562 cells using data from the ENCODE Consortium (1, 2) (Supplementary Figure 10A). Considering the substantial size of the output correlation matrix and the difficulties with visualization, we also set up a Shiny app (79) visualization tool for convenient and interactive exploration of the correlation matrix (Supplementary Figure 10B).

Lastly, we’d like to point out a potential limitation: JMnorm might not be able to adequately handle global changes across all functionally correlated features under certain conditions. This could result in losing some global changes at the within-cluster quantile normalization step. For these scenarios, other normalization strategies that take into account global changes, such as normalization using only common background regions (77) or a spike-in reference (75, 76), may be more appropriate.

In summary, JMnorm introduces a novel approach for multi-feature normalization of epigenetic data. This method has a straightforward design and is effective in reducing technical biases and preserving cross-feature correlations across different cell types. With the continuing development of high-throughput sequencing technologies for genome interrogation and the growing number of epigenetic datasets generated in different cell types, species, and under various physiological conditions, we anticipate JMnorm becoming a crucial tool for data normalization in integrative and comparative epigenomics studies.

## DATA AVAILABILITY

The JMnorm package is available at GitHub (https://github.com/camp4tx/JMnorm) (87) and released under GNU General Public License, version 2.0 or later. The main part of JMnorm was implemented in R. We also provided a conda environment that can be deployed in both MacOS and Linux operating systems. The signal tracks used in this project were mainly downloaded from VISION project data portal (32) and the EpiMAP repository (34). The detailed cell types included in each cell type group in the EpiMAP data can be found in the EpiMAP metadata file (88). Human and mouse cCRE lists were downloaded from ENCODE-SCREEN data portal (89, 90). The list of links for the files used in this paper can be found in Supplementary Table 1.

## ACKNOWLEDGEMENTS

We would like to express our gratitude to the ENCODE, VISION, and EpiMAP data Consortium projects for their valuable resources. We thank Gokul Ramaswami and Wei Jiang for their valuable suggestions and help.

