## Supplemental Material 1 for "JMnorm: a novel Joint Multi-feature normalization method for integrative and comparative epigenomics"

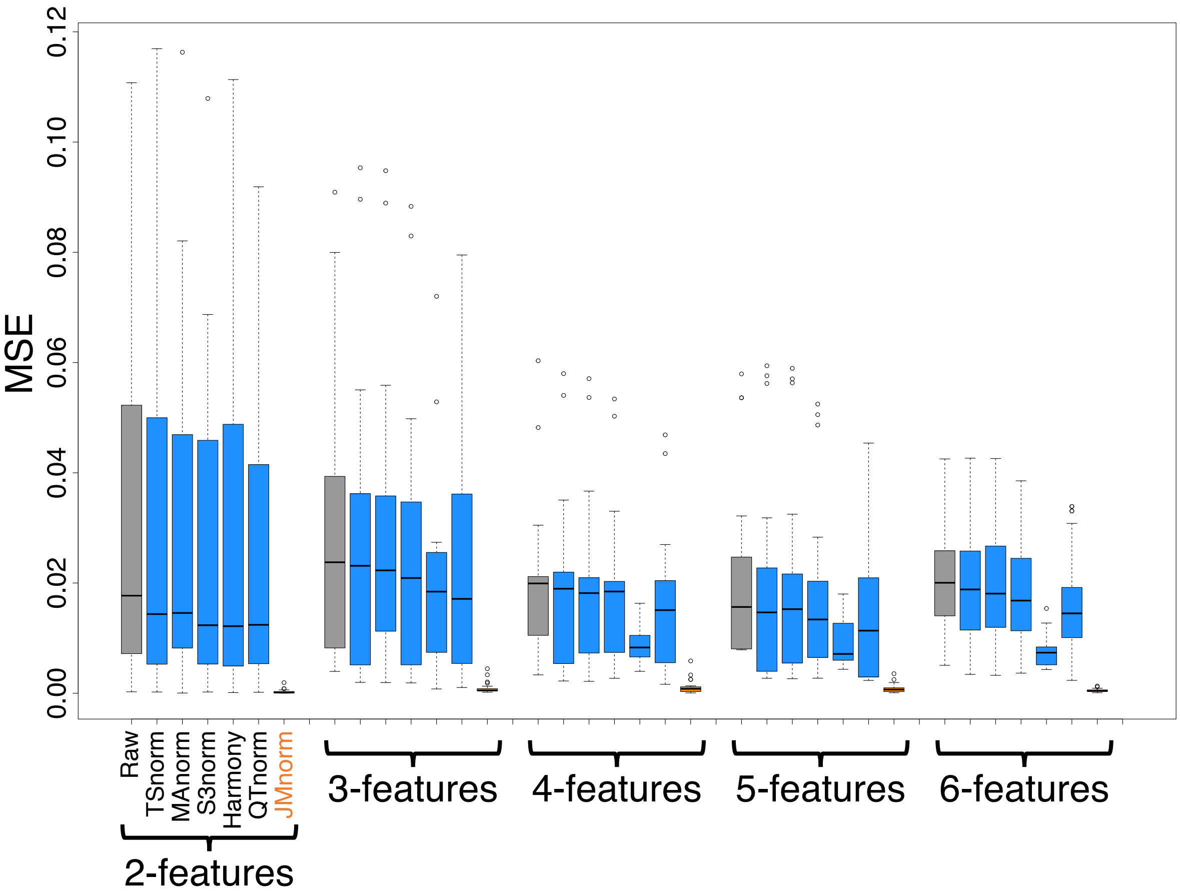


**Supplementary Figure 1.** A boxplot series of the mean squared errors (MSEs) between the target and reference cross-feature correlation matrices. The figure contains five sets of boxplots, with each set representing the results of normalization for data sets with different numbers of epigenetic features. The boxplots within each set represent the results for various methods, arranged in the following order: Raw signal (gray), TSnorm (blue), MAnorm (blue), S3norm (blue), Harmony (blue), QTnorm (blue), and JMnorm (orange).

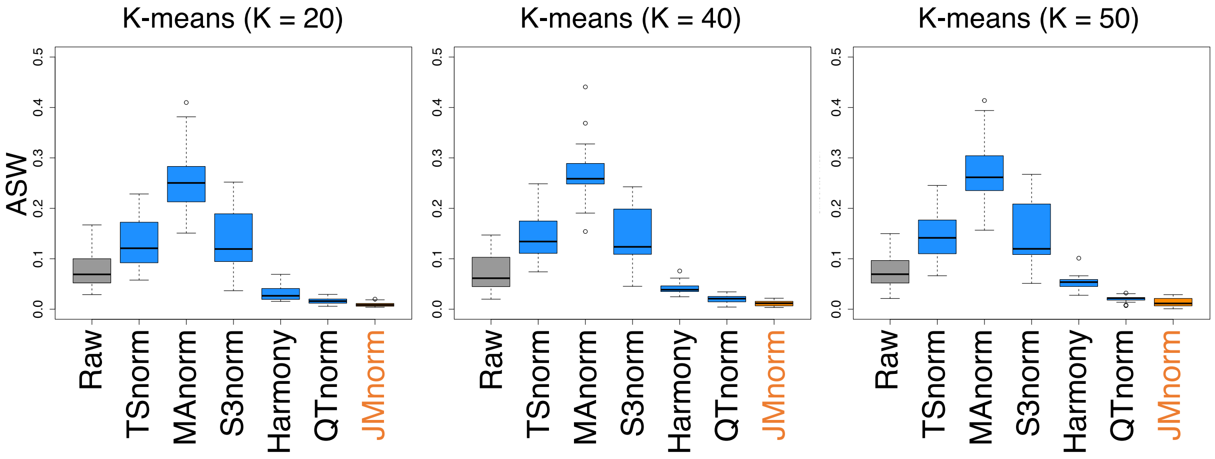


**Supplementary Figure 2.** A boxplot of average silhouette widths (ASWs) for the six normalization methods comparing quality of mixing of cCREs across different cell-types in clustering outputs (Kmeans: K=20, 40, 50). Lower ASW values indicate better mixing of cCREs across different cell types in clustering outputs.

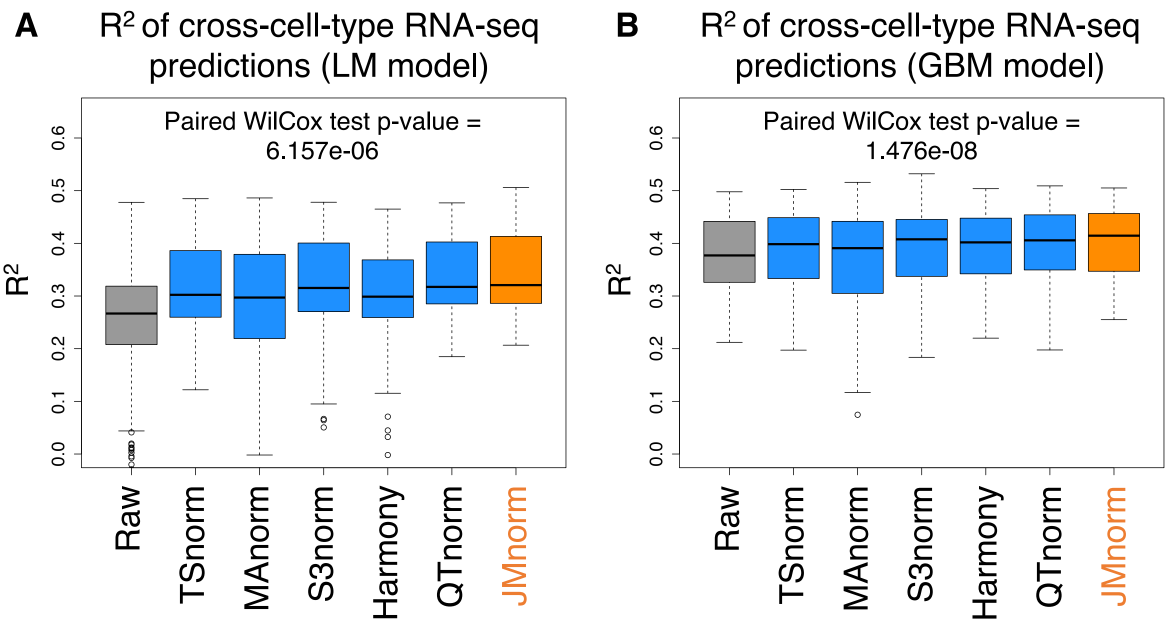


**Supplementary Figure 3.** Boxplots of $R^2$ values between observed and predicted RNA-seq log2TPMs for the six normalization methods using linear regression model (LM) (**Panel A**) and gradient boosting regression model (GBM) (**Panel B**) based on multi-feature signal matrices normalized using different methods. Higher $R^2$ values indicate better accuracy in cross-cell type hold-out genes' expression prediction.

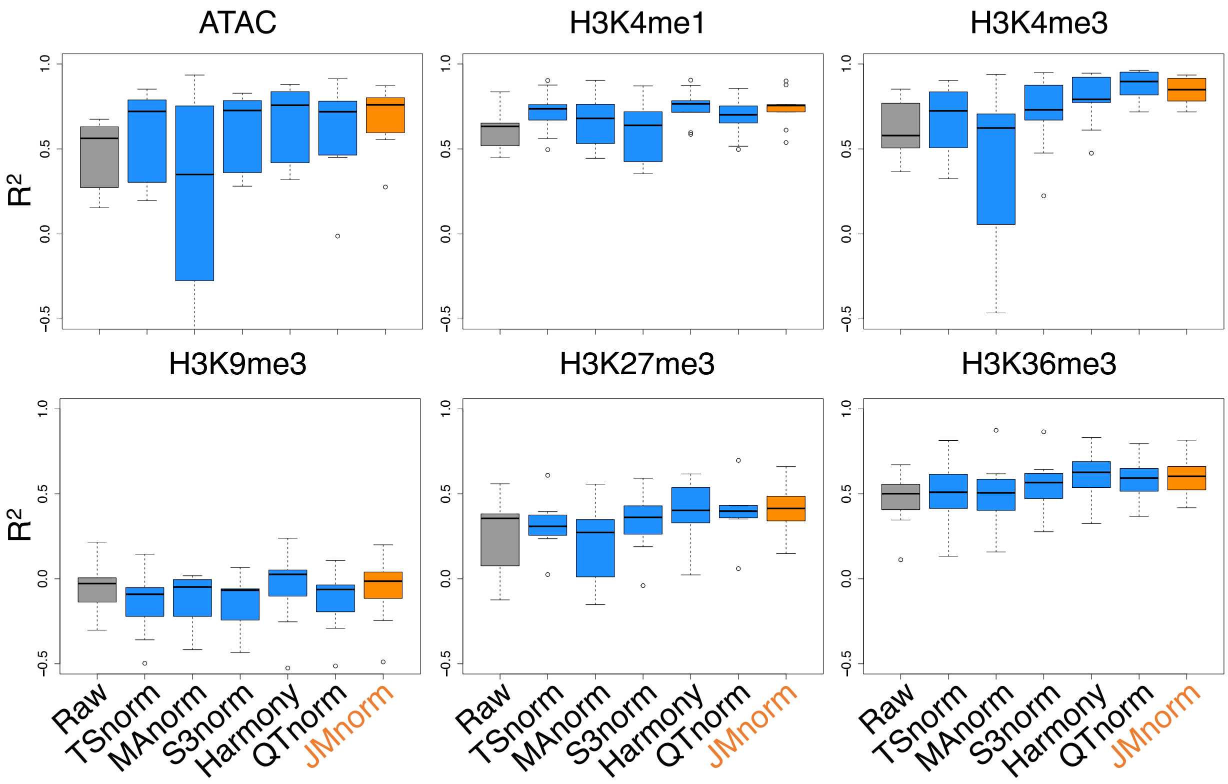


**Supplementary Figure 4.** The boxplots of $R^2$ values between biological replicates for the six normalization methods in multiple cell-types for six different epigenetic features.

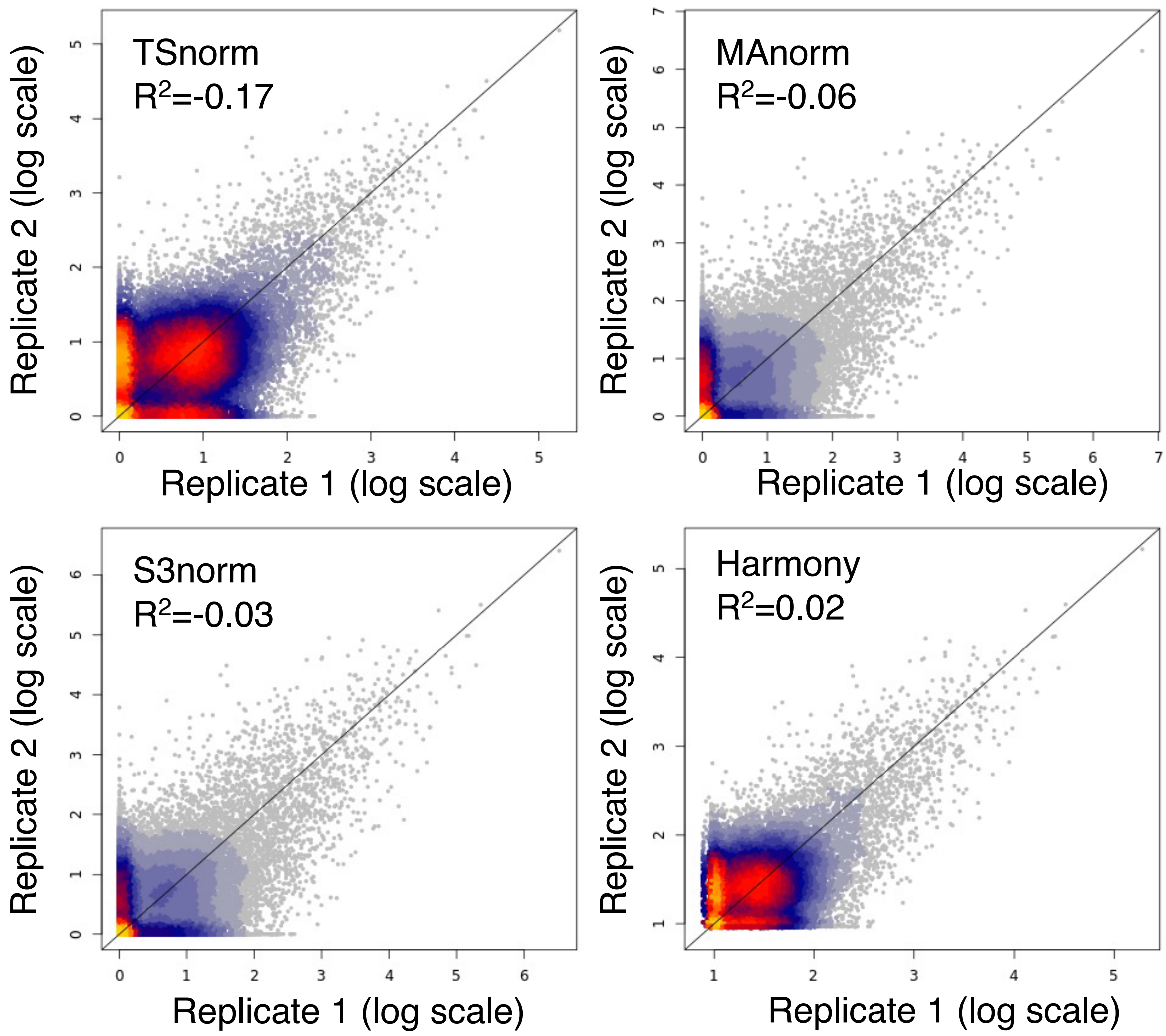


**Supplementary Figure 5.** Scatterplots of H3K27ac ChIP-seq signal in $CD8^{+}$ T-cells between two biological replicates normalized by four different methods (log2 scale). Bright orange and gray colors indicate higher and lower data point density, respectively.

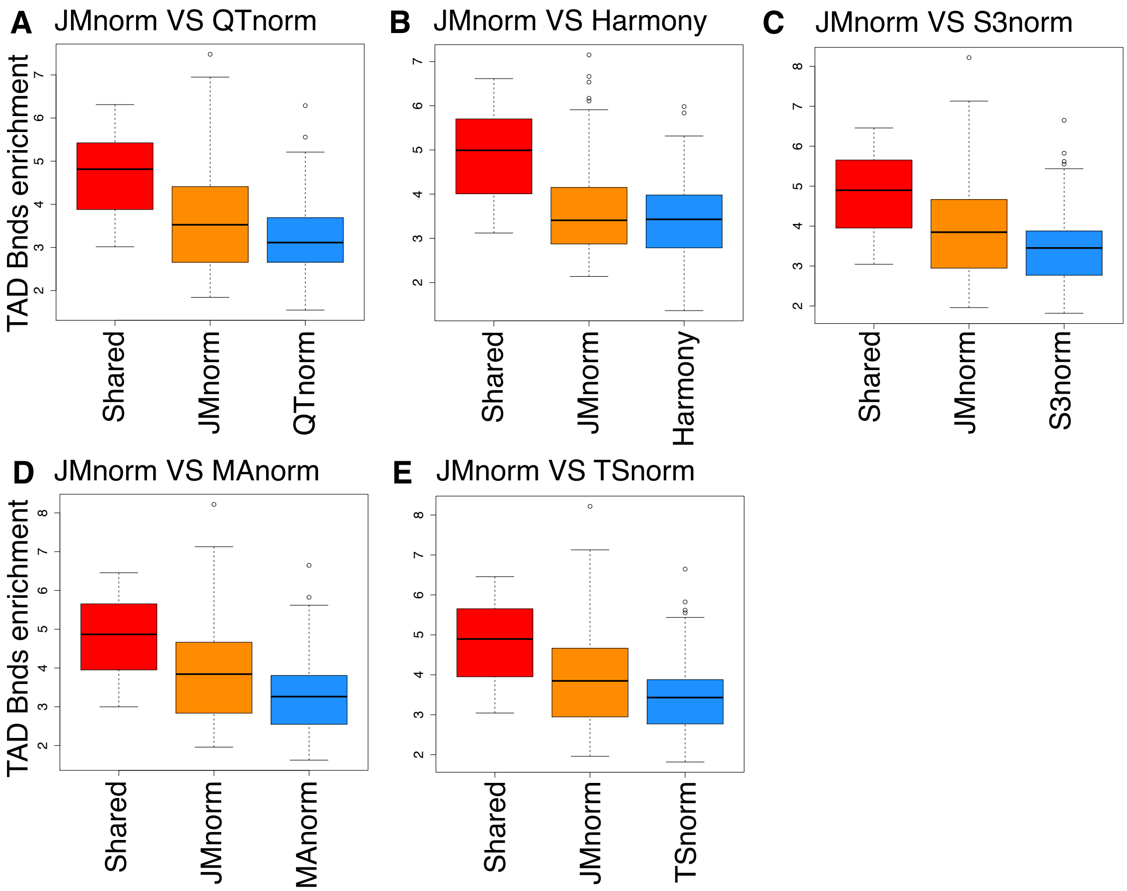


**Supplementary Figure 6.** Evaluation of quality of TF peak calling results. Box plots comparing JMnorm's and five other methods' performance as determined by CTCF peak enrichment at TAD boundaries. Red box plots represent enrichments for CTCF peaks that are shared between JMnorm and a corresponding alternative method. Orange box plots represent enrichments for CTCF peaks uniquely identified by JMnorm. Blue box plots represent enrichments for CTCF peaks uniquely identified by the respective alternative method.

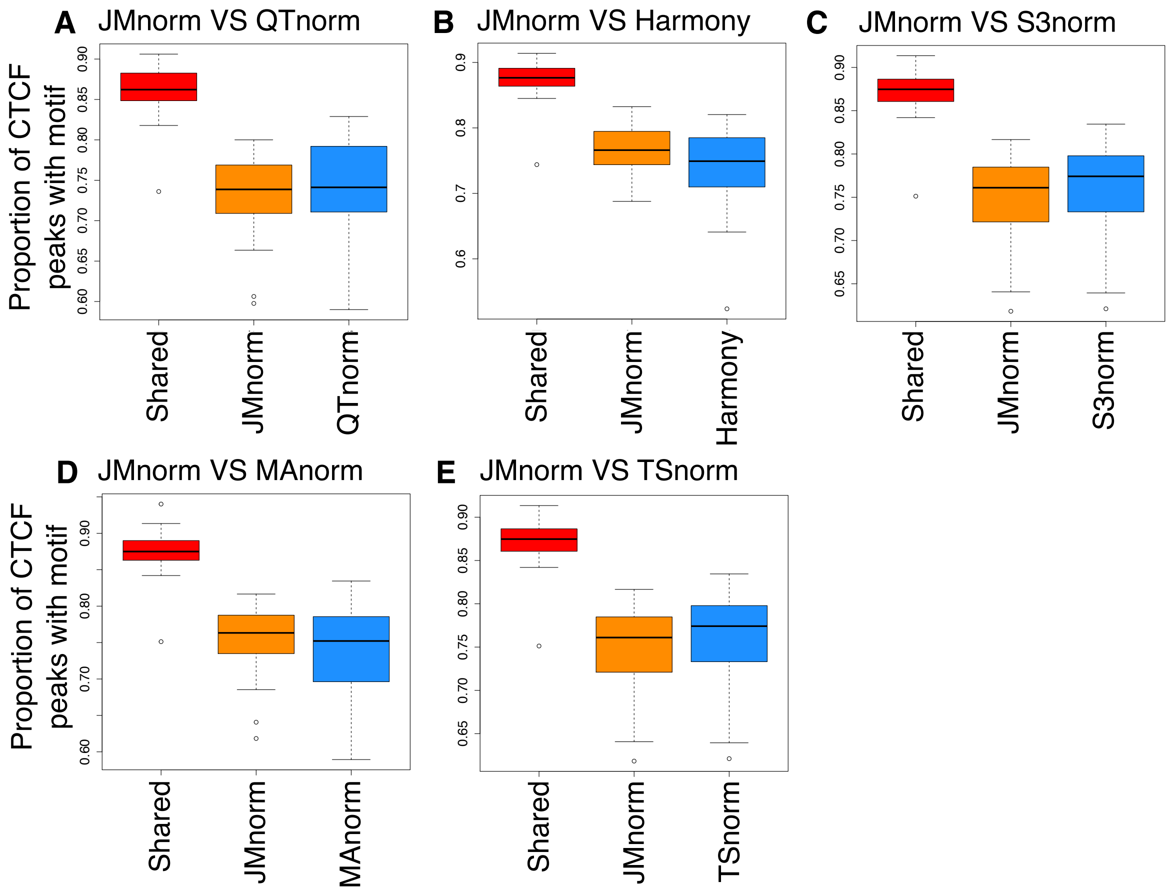


**Supplementary Figure 7.** Evaluation of quality of TF peak calling results. Box plots comparing JMnorm's and five other methods' performance as determined by the proportion of CTCF peaks with CTCF motif (Jaspar ID: MA0139.1). Red box plots represent proportions of CTCF peaks that are shared between JMnorm and a corresponding alternative method. Orange box plots represent proportions for CTCF peaks uniquely identified by JMnorm. Blue box plots represent proportions for CTCF peaks uniquely identified by the respective alternative method.

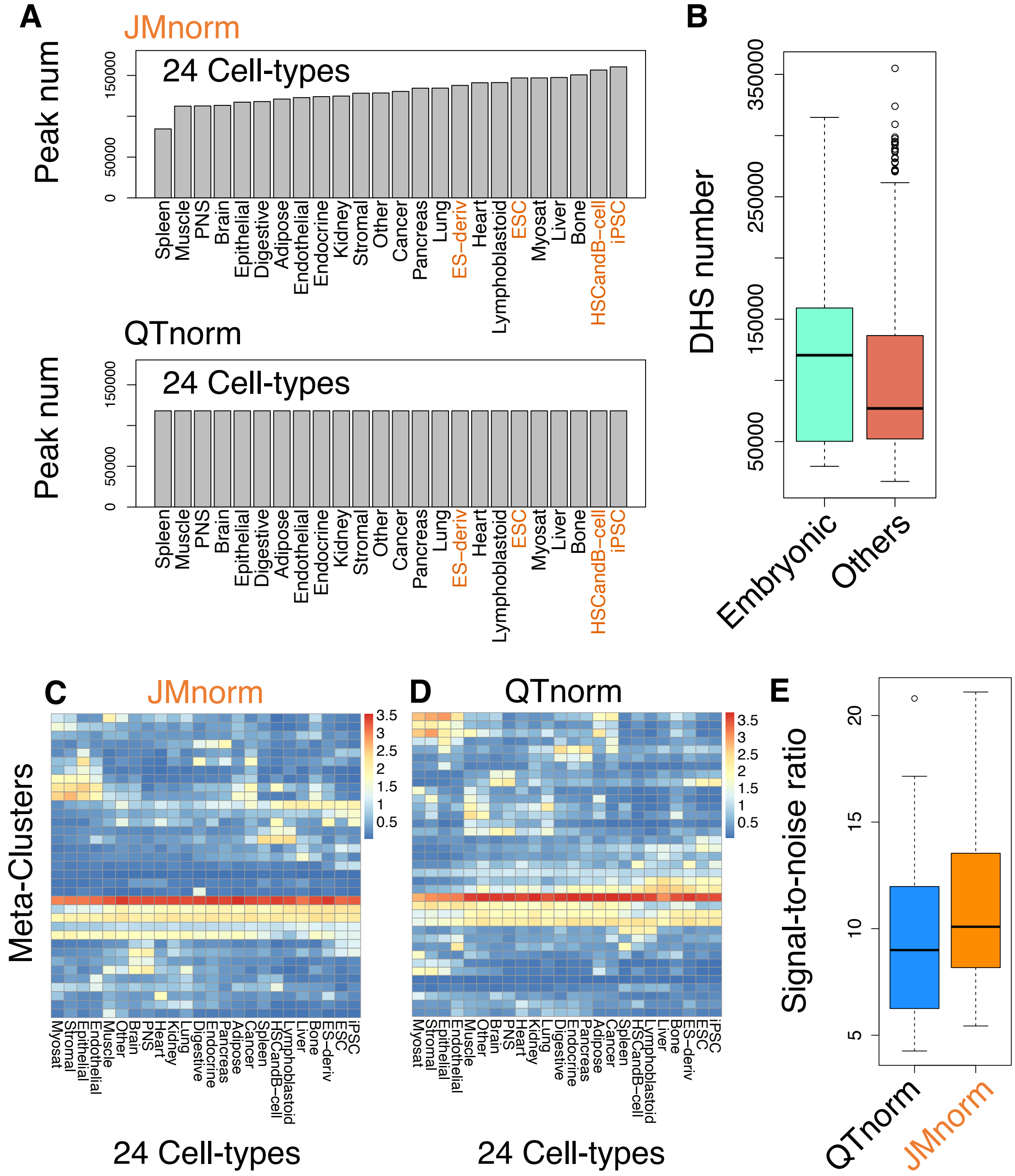


**Supplementary Figure 8.** Comparative analysis of JMnorm and QTnorm using DNase-seq data. (**A**) A boxplot comparing the number of DHSs identified in embryonic cell types with other cell types from the Meuleman et al. study. (**B**) Bar plots displaying the number of DNase-seq peaks detected using JMnorm-normalized data (top) and QTnorm-normalized data (bottom) across various cell types. Orange color highlights cell types that are closely related to stem cells. (C) Heatmaps of the average normalized DNase-seq signals for Snapshot Meta-Cluster generated by JMnorm. (D) same as (C), for QTnorm. (E) A boxplot of the signal-to-noise ratios for the Meta-Cluster average heatmaps.

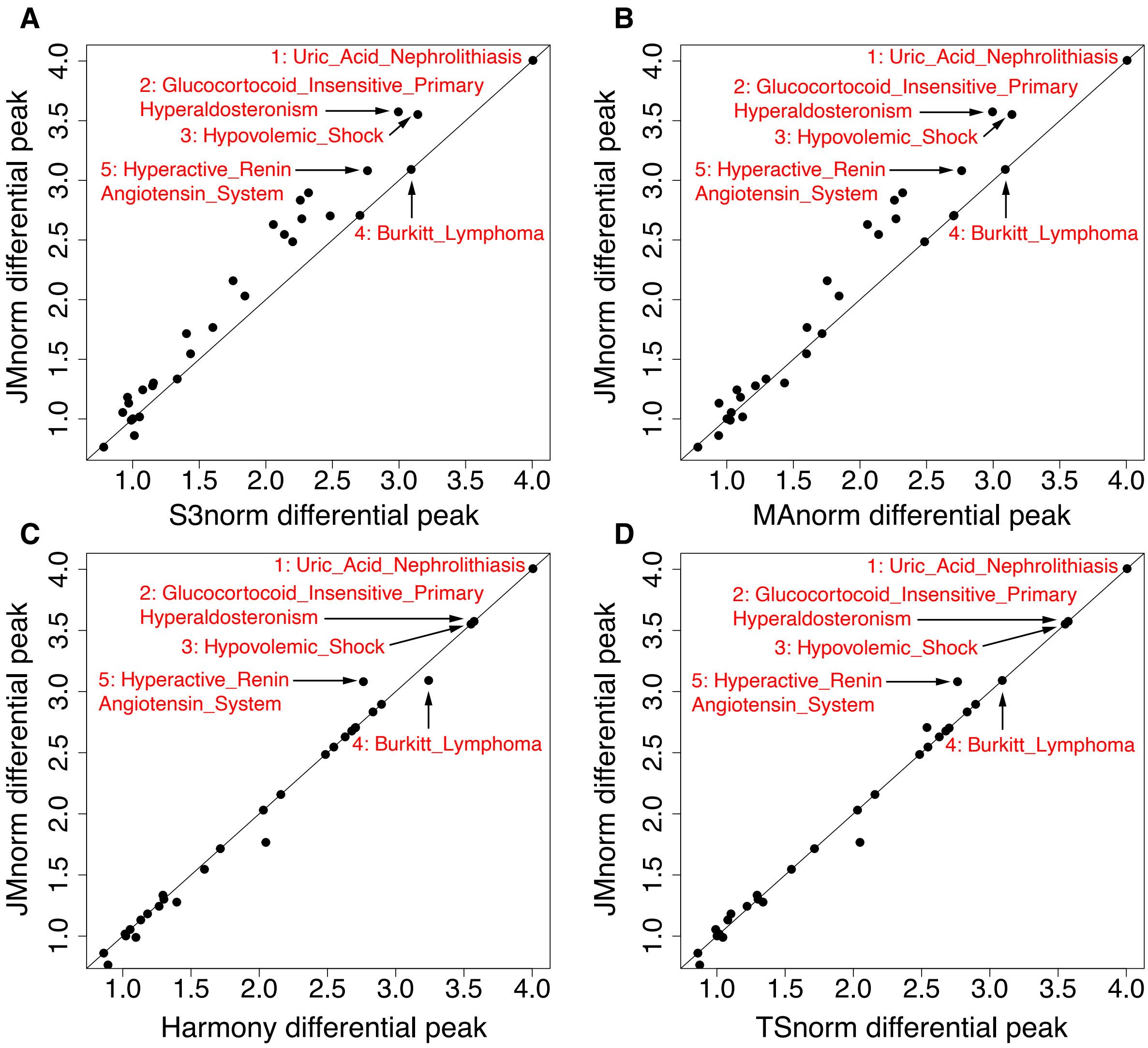


**Supplementary Figure 9.** Scatterplots of enrichments of the Human Phenotype terms in differential GR ChIP-seq peaks identified by JMnorm (y-axis) and S3norm (Panel A) / MAnorm (Panel B) / Harmony (Panel C) / TSnorm (Panel D) (x-axis).

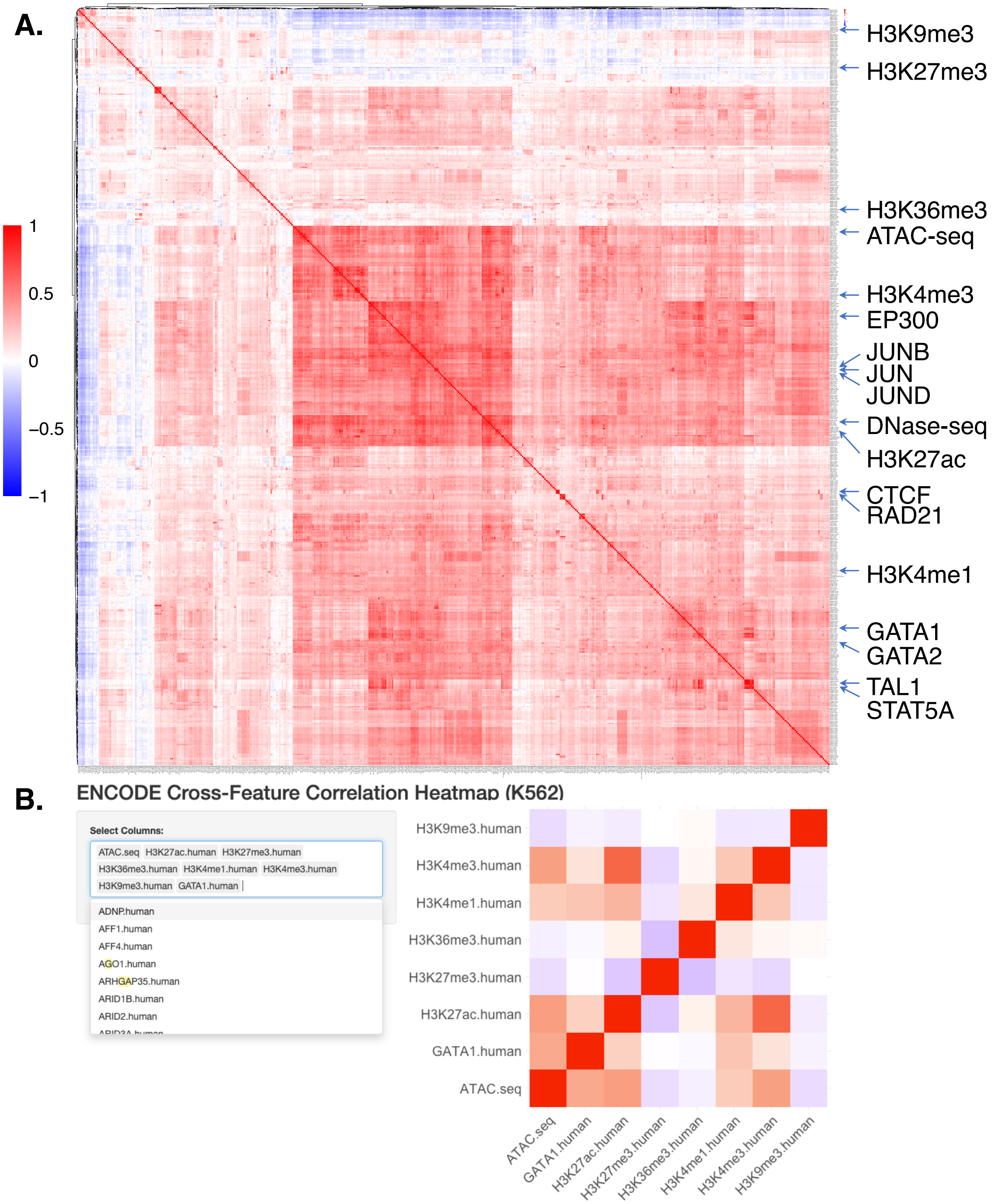


**Supplementary Figure 10.** (**A**) The Pearson correlation matrix of 538 epigenetic features in K562 cells. (**B**) The Shiny app visualization tool designed to aid in searching and visualization of a subset of features from the afore-mentioned correlation matrix.
